# Individual thalamic inhibitory interneurons are functionally specialized towards distinct visual features

**DOI:** 10.1101/2023.03.22.533751

**Authors:** Fiona E. Müllner, Botond Roska

## Abstract

Inhibitory interneurons in the dorsolateral geniculate nucleus (dLGN) are situated at the first central synapse of the image-forming visual pathway but little is known about their function. Given their anatomy, they are expected to be multiplexors, integrating many different retinal channels along their dendrites. Here, using targeted single-cell-initiated rabies tracing, we found that mouse dLGN interneurons exhibit a degree of retinal input specialization similar to thalamocortical neurons. Some are anatomically highly specialized, for example, towards direction-selective information. Two-photon calcium imaging performed *in vivo* revealed that interneurons are also functionally specialized. In mice lacking retinal horizontal direction selectivity, horizontal direction selectivity is reduced in interneurons, suggesting a causal link between input and functional specialization. Functional specialization is not only present at interneuron somata, but also extends into their dendrites. Altogether, each inhibitory interneuron globally encodes one visual feature originating mostly in the retina and is ideally suited to perform feature-selective inhibition.

## INTRODUCTION

All sensory systems, except for smell, pass through the thalamus as information flows from the sensory periphery to the cortex. How sensory information is modified within the thalamus and the function of the thalamus in sensory processing are key and open questions in neuroscience. The dorsolateral geniculate nucleus (dLGN) is the primary visual thalamic nucleus necessary for image perception. Retinal ganglion cells, as the output neurons of the retina, synapse onto excitatory thalamocortical neurons in the dLGN, which in turn project to the primary visual cortex. The dLGN also contains GABAergic inhibitory interneurons that receive inputs from retinal ganglion cells and synapse onto thalamocortical neurons (Sherman, 2004). The interneurons are ideally suited to influence visual information at this early stage (Hirsch et al., 2015; Wang et al., 2011) but little is known about their *in vivo* function or how they process retinal information.

The retina of vertebrates extracts many features from the visual scene and sends these features to the dLGN via the axons of distinct ganglion cell types. Mice have 46 transcriptomic ganglion cell types (Tran et al., 2019), each of which has characteristic morphological and electrophysiological properties (Goetz et al., 2022). Which visual feature a specific ganglion cell type extracts (the “retinal channel”) is determined by the specific bipolar and amacrine cell input it receives (Roska and Werblin, 2001) and its electrophysiological properties (Wienbar and Schwartz, 2022). The axons of bipolar cells and the processes of amacrine cells that provide input to the dendrites of ganglion cells are organized in layers within the inner plexiform layer of the retina (Wässle, 2004). Accordingly, the dendrites of each ganglion cell type display a stereotypical stratification pattern within the inner plexiform layer. Neuronal processes of starburst amacrine cells, which express choline acetyltransferase (ChAT), form two distinct layers that can serve as anatomical landmarks (ChAT-bands). Relative to these two bands, the inner plexiform layer can be divided into ten strata (Siegert et al., 2009). Functional order exists across these strata: the dendrites of ganglion cells that respond to increases in visual intensity (ON cells) reside in strata 6-10, while dendrites of ganglion cells that respond to decreases in visual intensity (OFF cells) are found in strata 1-4. Ganglion cells with dendrites in both divisions respond to both increases and decreases of visual intensities (ON-OFF cells, Masland, 2012). Ganglion cells with dendrites in the outermost strata 1, 2 and 8-10 show more sustained responses to light increments or decrements, whereas those with dendrites in the innermost strata 3-7 show more transient responses (Baden et al., 2013; Goetz et al., 2022; Roska and Werblin, 2001). Direction-selective retinal ganglion cells co-fasciculate with the processes of starburst amacrine cells and, therefore, co-stratify within the ChAT-positive layers 3 and 7.

Within the dLGN, retinal information was long considered to stay in “labeled lines”, with each thalamocortical neuron receiving information from one type of retinal ganglion cell (Cleland et al., 1971; Mastronarde, 1992; Usrey et al., 1999). Recent evidence suggests more complex and even binocular visual processing (Hammer et al., 2015; Howarth et al., 2014; Litvina and Chen, 2017; Morgan et al., 2016; Román Rosón et al., 2019; Rompani et al., 2017; Zeater et al., 2015). In mice, only 28% of thalamocortical neurons combine retinal inputs from one ganglion cell type and can thus be considered as labeled lines (“relay-mode”), while the remaining thalamocortical neurons combine information from different retinal ganglion cell types, either from the contralateral eye (“combination mode”) or from both eyes (“binocular mode”, Rompani et al., 2017).

GABAergic interneurons in the dLGN differ from thalamocortical neurons but also from cortical interneurons, most prominently by not only receiving synapses on their dendrites but also carrying output synapses on their dendrites. These unconventional dendritic outputs are the most prevalent output synapses of dLGN interneurons, the so-called F2 synapses (Colonnier and Guillery, 1964; Guillery, 1969; Szentágothai et al., 1966). Dendro-dendritic F2 synapses are often found in a triadic arrangement, i.e. the same retinal ganglion cell axon synapses onto both the interneuron dendrite and the thalamocortical neuron dendrite, to which the interneuron provides a GABAergic inhibitory synapse in the immediate vicinity. Triadic output synapses have been found in the thalamus of many different mammalian species (Sherman, 2004). In a 3D volume of dLGN consisting of a fully reconstructed mouse dLGN interneuron, 94% of retinal ganglion cell axonal boutons that contacted the dLGN interneuron also contacted a thalamocortical neuron dendrite, and 90% of these retinal input synapses were found in a triadic motif, such that the interneuron provided input to the same thalamocortical dendrite as the retinal axon (Morgan and Lichtman, 2020).

In addition to having dendritic output synapses, it was suggested that the cable properties of the dLGN interneurons yield strong attenuation of signals along their dendrites (Bloomfield et al., 1987; Bloomfield and Sherman, 1989). The abundance of dendro-dendritic output synapses, together with the predicted strong attenuation of their branched dendritic arbor, have given rise to the hypothesis that dendrites of dLGN interneurons act as ‘Multiplexor’ devices, with many independent processing units (Bloomfield and Sherman, 1989; Cox et al., 1998; Cox and Sherman, 2000; Govindaiah and Cox, 2006; Pare et al., 1991).

Functionally, the triadic output is thought to provide fast feedforward inhibition that shapes the incoming retinal information in the time domain. This fast inhibition was suggested to provide contrast gain control (Sherman, 2004), remove secondary spikes (Blitz and Regehr, 2005), or introduce a lag in responses that could be used to compute direction selectivity in cortex (Blitz and Regehr, 2005; Vigeland et al., 2013). The dendrites of rodent dLGN interneurons span large parts of the dLGN (Morgan and Lichtman, 2020; Zhu et al., 1999) and lack the age-related pruning observed in thalamocortical neurons (Seabrook et al., 2013) that was suggested to contribute to the specialization of thalamocortical neurons towards selected retinal ganglion cell types. Taken together, these properties lead to the prediction that dLGN interneurons sample retinal information broadly and unselectively along their local and multiplexing inhibitory input-output units.

In contrast to their supposed role as ‘Multiplexors’, global calcium spikes as well as action potentials have been observed in dLGN interneurons *in vitro* (Casale and McCormick, 2011), which points to the possibility that they might perform cell-wide signaling. We therefore asked whether dLGN interneurons indeed sample retinal information unselectively, as expected if their dendritic units are independent, or whether they exhibit preference towards specific retinal ganglion cell types. Preference towards specific retinal inputs could result in visual response selectivity of the interneurons and contribute to a cell-wide signaling function. To investigate this, we developed an approach to image LGN interneurons *in vivo* and asked how diversely interneurons respond to visual stimuli and whether their responses are causally related to the visual features of their retinal ganglion cell inputs.

Our results suggest that, instead of being ‘Multiplexors’, dLGN interneurons act as ‘Selectors’, receiving inputs from a defined set of retinal ganglion cell types that cause them to selectively represent specific visual features. Individual interneurons represent the same feature cell wide, while different interneurons exhibit diverse features that form a continuum in the feature space. The anatomical and functional properties of feature-selective dLGN interneurons make them ideal candidates to mediate feature-selective inhibition at the first central synapse of image-forming vision.

## RESULTS

### *In vivo* single-cell-initiated monosynaptic rabies tracing from dLGN interneurons

To determine the number and types of retinal ganglion cells that provide synaptic inputs to individual dLGN inhibitory interneurons, we developed a strategy to perform *in vivo* single-cell-initiated monosynaptic rabies tracing (Wickersham et al., 2007) from genetically identified mouse dLGN interneurons. We labeled dLGN interneurons fluorescently using transgenic GAD-lines − GAD65-IRES-cre (Taniguchi et al., 2011) crossed to EYFP-reporter (Ai3, Madisen et al., 2010) or GAD67-GFP (Tamamaki et al., 2003) lines − and targeted the labeled neurons by two-photon imaging for *in vivo* single-cell electroporation (Figure 1A). In each mouse, we electroporated a dLGN interneuron with fluorescent Alexa-594 dye, to monitor the success of the electroporation (Figure 1B), together with three plasmids (Rompani et al., 2017; Wickersham et al., 2007): the avian TVA receptor, the rabies G glycoprotein, and the fluorescent protein tdTomato. Finally, we injected a G-deleted (SADΔG), EnvA-coated rabies virus expressing mCherry into the dLGN. EnvA, the ligand of TVA, permits entry of the rabies virus exclusively into the TVA-expressing cell (Figure 1C), and the G glycoprotein enables transsynaptic transfer of rabies virus to presynaptic partners of the targeted dLGN interneuron (Wickersham et al., 2007).

**Figure 1:**
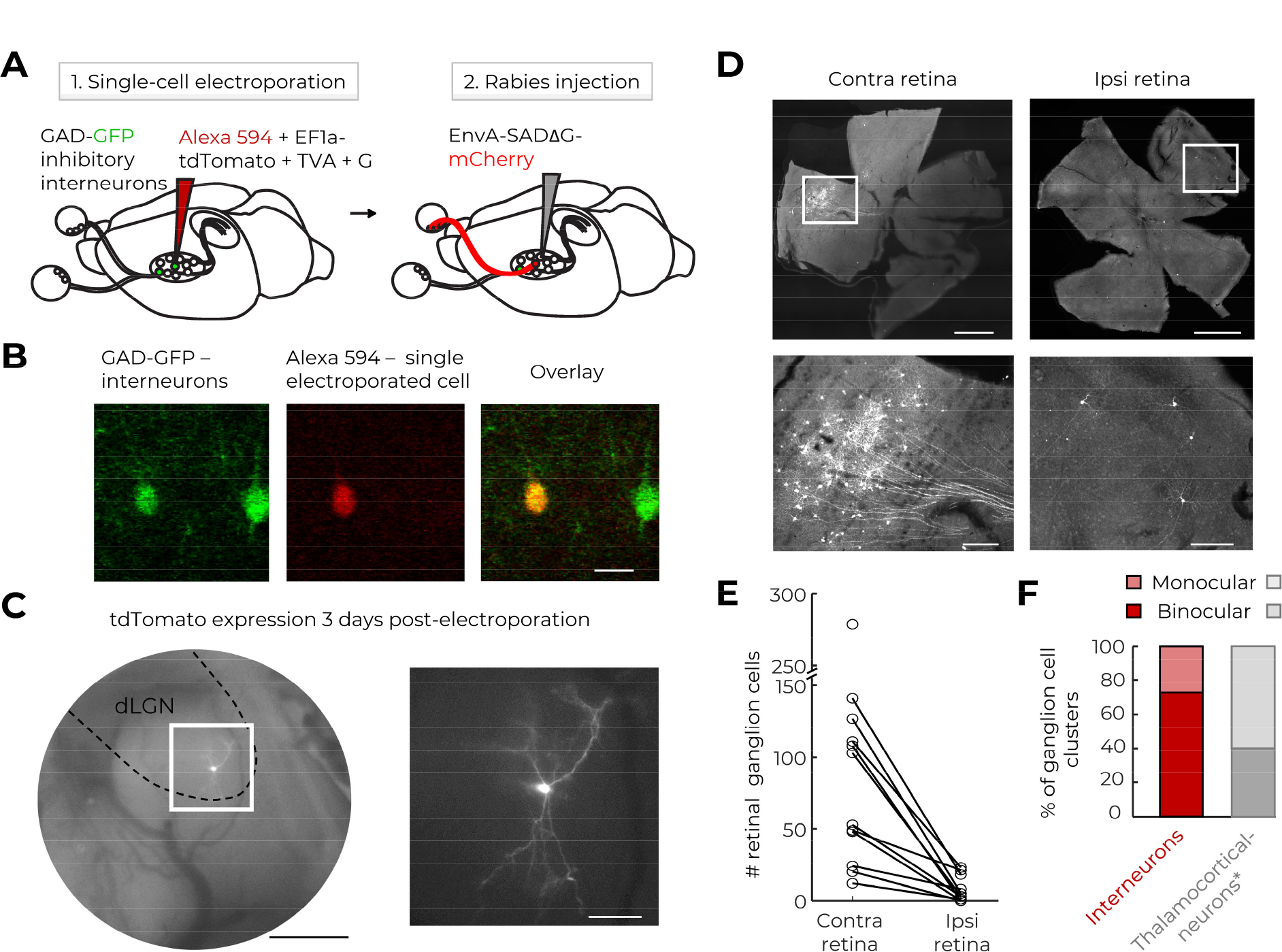
Single-cell-initiated rabies tracing from genetically identified deep-brain interneurons. **A:** Single-cell-initiated rabies tracing: First, tdTomato, TVA, and G are delivered to a GFP-labeled interneuron via electroporation. Second, EnvA-SADΔG-mCherry rabies virus enters the TVA-G-expressing cell and spreads retrogradely from the infected cell. **B:** Two-photon imaging-guided targeted single-cell electroporation of a GAD-positive dLGN interneuron (green) results in Alexa 594 filling (red) of the electroporated neuron but not neighboring neurons. Scalebar: 20 µm. **C:** After 3 days of incubation, the expression of tdTomato is monitored through the cranial window (left, scalebar: 500 µm). Zoom in: electroporated interneuron displaying its characteristic morphology (scalebar: 100 µm). **D:** Presynaptic retinal ganglion cells in contra- and ipsilateral retinas that project to a single dLGN interneuron. Upper panels: low magnification, scalebars: 1000 µm. Lower panels: zoom in, scalebars: 200 µm. **E:** Presynaptic retinal ganglion cell counts in contra- and ipsilateral retinas. **F:** Percentage of monocular and binocular presynaptic retinal ganglion cell clusters. Data from thalamocortical neurons were reported previously (Rompani et al., 2017).

With this approach, we mapped presynaptic retinal ganglion cell inputs to 12 dLGN interneurons (Figure 1D). As expected from their large dendritic size (Figure 1C; Morgan and Lichtman, 2020; Zhu et al., 1999), dLGN interneurons received inputs from a greater number of retinal ganglion cells than thalamocortical neurons (mean of 89.6 ganglion cells contralateral and 7.5 ipsilateral compared to 15.2 ganglion cells contralateral and 12.6 ipsilateral in thalamocortical neurons; Figure 1E and Supplemental Figure 1, thalamocortical neuron data from Rompani et al., 2017). Also consistent with their wide dendritic arbor, a large proportion (72.7%) of interneurons received inputs from both eyes (Figure 1F). All binocular interneurons received more inputs from the contralateral than the ipsilateral retina, with an average contra/ipsi ratio of 16.3. In contrast to that observed for thalamocortical neurons, the distribution of the number of ganglion cells in the contra- and ipsilateral retina providing input to interneurons did not deviate significantly from a random- binomial distribution (p=0.66 versus p=0.004 for interneurons versus thalamocortical neurons).

This was consistent with the initial hypothesis that interneurons sample retinal information randomly.

To understand which ganglion cell types provide inputs to dLGN interneurons, we classified the presynaptic retinal ganglion cells based on the stratification of their dendrites within the inner plexiform layer (Figure 2A). We took high-resolution confocal stacks of the presynaptic ganglion cells in flat-mounted retinas co-stained with antibodies against mCherry and ChAT (Figure 2B). Previous analysis of the rabies-traced ganglion cells (Rompani et al., 2017) had two bottlenecks. First, the assignment of dendrites to individual cells required manual segmentation. Second, the stratification of dendrites within the inner plexiform layer was assessed based on successive side projections of the segmented dendrites, which was also performed manually. Both bottlenecks arise from the three-dimensional nature of the flat-mounted retinas, in which the ChAT-bands are curved planes (Figure 2B). To overcome these limitations, we trained a convolutional neural network, U-Net, to automatically detect the ChAT-bands. Knowing the ChAT-band locations in the z-axis, we could virtually flatten the image stacks (Figure 2B). The dendrites of mCherry-labeled ganglion cells could then, in a single top-down maximum projection, be automatically labeled with different colors according to their depths relative to the ChAT-bands (Figure 2C). In this pseudo-colored top-down projection, dendrites belonging to a given ganglion cell can be recognized by their emergence from the soma in a snowflake-crystal pattern, and the dendrites can be followed traveling through the layers of the inner plexiform layer as their color changes in a rainbow sequence (Figure 2C, Supplemental Figure 2). For binocular cells, we determined the retinal ganglion cell types separately in both eyes, and we refer to the presynaptic ganglion cells in each retina as “cluster”, since the ganglion cells were typically centered in one quadrant (Figure 1D). We classified 638 retinal ganglion cells labeled in single-cell-initiated rabies tracings from dLGN interneurons.

**Figure 2:**
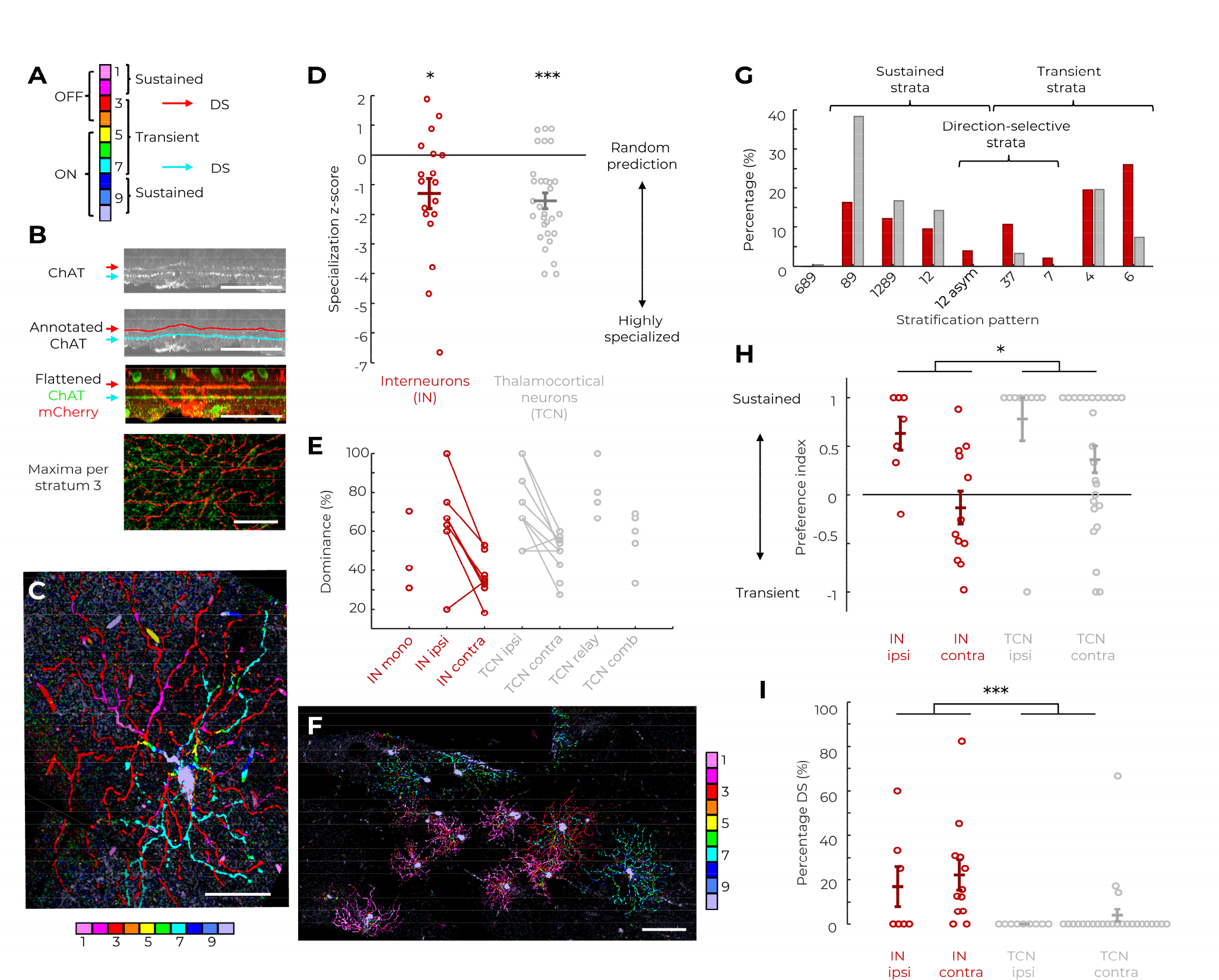
Retinal ganglion cell inputs to individual dLGN interneurons are specialized. **A:** Correlation between stratification and function for ganglion cell dendrites. Numbers refer to strata of the inner plexiform layer. DS: direction selective. **B:** Machine-learning assisted detection of the ChAT-bands and artificial flattening of the retina. Panels from top to bottom show: ChAT-stained retina with curvature; annotated ChAT-bands (red and cyan); flattened z-stack with ChAT-bands (green) and rabies-infected ganglion cells (red); maximum z-projection through stratum 3 visualizes co-fasciculation of the ganglion cell dendrites and ChAT-positive processes. Scalebars: 30 µm. **C:** A maximum z-projection of the example ganglion cell in **B** is pseudo-colored based on relative distance of the maxima from the ChAT-bands. In this example, the dendritic stratification (cyan/red) of an ON-OFF direction-selective retinal ganglion cell is visible. Scalebar: 30 µm. **D:** Specialization z-scores for all retinas with ganglion cells presynaptic to dLGN interneurons (red) or thalamocortical neurons (gray). *: p=0.020, ***: p<0.001, Wilcoxon test against zero with Bonferroni-Holm correction. **E:** Dominance values for interneurons (IN, red) and thalamocortical neurons (TCN, gray). Values for individual retinal clusters are grouped as follows. Mono: monocular, ipsi: ipsilateral, contra: contralateral, relay: relay-mode, comb: combination-mode. Relay- and combination-mode refer to the definitions in Rompani et al., 2017. **F:** Example of a contralateral ganglion cell cluster (zoom-in to cluster center) consisting of JAM- B (pink), ON-OFF direction-selective (cyan + red) and ON direction-selective (cyan) retinal ganglion cells. Scalebar: 100 µm. **G:** Histogram of stratification patterns of presynaptic ganglion cells transsynaptically labeled from interneurons (red) or thalamocortical neurons (gray). Strata “xyz” refers to ganglion cells stratifying in retinal strata x, y, and z. **H:** Preference indices shown for ganglion cell clusters found in ipsi- and contralateral retinas labeled from interneurons (IN, red) and thalamocortical neurons (TCN, gray). *: p=0.039, Mann- Whitney-U test. **I:** The percentage of direction-selective inputs to interneurons (IN, red) and thalamocortical neurons (TCN, gray) (co-stratification with strata 3, 7 or stratum 7, or stratification in strata 1, 2 with asymmetric dendrites characteristic of JAM-B cells) from ipsi- and contralateral retinas. ***: p<0.001, Mann-Whitney-U test. **D,E,G,H,I**: *n*=638 ganglion cells presynaptic to 12 interneurons; *n*=245 ganglion cells presynaptic to 15 thalamocortical neurons. Horizontal lines indicate group averages, error bars indicate ± sem. All thalamocortical neuron data were taken from Rompani et al., 2017.

### Retinal ganglion cell inputs to dLGN interneurons are specialized

We used two approaches to quantify the specialization of retinal ganglion cell inputs to dLGN interneurons. First, we measured the difference in the number of ganglion cell types presynaptic to each dLGN interneuron from that expected by a random draw. For each number of presynaptic ganglion cells, we simulated a random draw from the empirically found overall distribution of ganglion cell types using Monte-Carlo simulation. The number of ganglion cell types expected from a random draw depends on the total number of retinal ganglion cells (Supplemental Figure 2). The deviation of the empirically found number of ganglion cell types to the number of ganglion cell types expected by a random draw was quantified by a specialization z-score (measured minus expected number, divided by the simulated standard deviation). A z-score of zero means that the inputs match a random prediction, and a negative z-score indicates that the inputs are specialized towards certain ganglion cell types. Given the Multiplexor model, we expected the dLGN interneurons to randomly sample retinal ganglion cell types and, therefore, to have z-scores around zero. In contrast, we found the z-scores of interneurons and thalamocortical neurons to be similarly distributed – with some more randomly sampling cells around zero and several highly specialized cells with z-scores smaller than -2 (Figure 2D, Supplemental Figure 2, both medians negative, Wilcoxon test, interneurons: p=0.020 and thalamocortical neurons: p<0.001).

Second, we analyzed interneuron input specialization by quantifying ganglion-cell-type dominance. Within a presynaptic cluster of retinal ganglion cells, ganglion cell types can be present in similar numbers, or the cluster can be dominated by an individual ganglion cell type. We defined dominance as the relative abundance of the most prevalent ganglion cell type, such that clusters dominated by individual ganglion cell types would have dominance values close to 100%, whereas clusters with equal representation of cell types would have lower dominance (e.g., 20% for five cell types). In analogy to thalamocortical neurons (TCN, Rompani et al., 2017), in which we had found monocular relay-mode and combination-mode clusters, as well as ipsi- and contralateral binocular clusters (‘TCN relay’, ‘TCN comb’, ‘TCN ipsi’, and ‘TCN contra’), we differentiated here for interneurons (IN) between monocular, binocular ipsilateral and binocular contralateral clusters (‘IN mono’, ‘IN ipsi’, ‘IN contra’). The dominance of presynaptic retinal ganglion cell clusters was quantitatively comparable between interneurons and thalamocortical neurons (Figure 2E): Similar dominance values were found in interneurons and thalamocortical neurons for both ipsilateral clusters (mean 65% and 73%, Mann-Whitney-U test, p=0.49) and contralateral clusters (mean 37% and 48%, Mann-Whitney-U test, p=0.082). Also, monocular presynaptic clusters of interneurons (‘IN mono’) and monocular combination-mode clusters of thalamocortical neurons (‘TCN comb’) exhibited similar dominance values (Mann-Whitney-U test, p=0.88), consistent with the fact that we observed no interneurons receiving relay-mode inputs formed by only one ganglion cell type. Even highly specialized interneurons received inputs from more than one ganglion cell type (Figure 2F). Most (86%) binocular interneurons showed higher dominance values for ipsilateral than contralateral clusters similar to those observed for binocular thalamocortical neurons (78%, mean difference contra-ipsi 28% (IN) and 24% (TCN), Mann-Whitney-U test, p=0.61).

While the degrees of specialization between LGN interneurons and thalamocortical neurons were similar, the sources of specialization were complementary (Figure 2G). We anatomically classified the retinal ganglion cells based on their stratification as putatively direction selective and putatively transient or sustained by the following rules: We defined retinal ganglion cells co-stratifying with the ChAT-bands (type 37 and 7) as well as ganglion cells stratifying in strata 1, 2 and having asymmetric dendritic arbors, characteristic of JAM-B cells (Kim et al., 2008), as direction selective. Similarly, we defined ganglion cells with dendrites in the innermost strata 3-7 as transient, while those in the outermost strata 1, 2 and 8-10 as sustained (Baden et al., 2016, 2013; Bae et al., 2018; Goetz et al., 2022; Roska and Werblin, 2001). Interneurons received fewer sustained and more transient and direction-selective retinal ganglion cell inputs than thalamocortical neurons (Supplemental Figure 2). 58% of retinal inputs to interneurons, compared to 30% to thalamocortical neurons, were transient (Fisher’s exact test, p<0.001). The percentage of direction-selective inputs was 15.1% for interneurons, as opposed to 3.3% for thalamocortical neurons (Fisher’s exact test, p<0.001). The percentage of interneurons receiving at least one direction-selective ganglion cell input (83%) was also significantly higher than for thalamocortical neurons (12%, Fisher’s exact test, p<0.001). The preference for transient inputs was especially pronounced in the contralateral inputs (61.8%). Retinal ganglion cell inputs from the ipsilateral retina, on the other hand, showed a preference towards sustained inputs, similar to the ipsilateral inputs to thalamocortical neurons, albeit with a slightly higher relative contribution of transient inputs (13% versus 3.6%). We calculated preference indices for individual retinas (Figure 2H) defined as the difference between retinal ganglion cell numbers divided by their sum: (# sustained - # transient) / (# sustained + # transient). A preference index of 1 indicates exclusively sustained, a preference index of -1 exclusively transient ganglion cells in a cluster. Preference indices spread widely, but the majority (58%) of contralateral retinas showed a preference for transient information, while 86% of ipsilateral retinas had preference for sustained information and 43% even received exclusively sustained information (preference index = +1). The preference indices for retinal clusters presynaptic to interneurons were significantly shifted towards transient information compared to that of thalamocortical neurons (Mann-Whitney-U test, p=0.039, Figure 2H, Supplemental Figure 2). Also, the percentages of direction selective inputs were significantly higher to interneurons than to thalamocortical neurons (Mann-Whitney-U test, p<0.001, Figure 2I, example shown in 2F, Supplemental Figure 2). Taken together, dLGN interneurons displayed a wide range of input specializations. Although the degrees of specialization resembled those of thalamocortical relay neurons, they displayed an overrepresentation of transient information and a significantly larger proportion of direction-selective inputs.

### dLGN interneurons are functionally specialized *in vivo*

The specialization of the ganglion cell types providing input to individual interneurons suggests that interneurons exhibit a range of different functional specializations. To test this, we established *in vivo* recordings from mouse interneurons by applying two-photon calcium imaging in a mouse line in which GCaMP6s-expression was limited to interneurons (GAD-IRES-cre, Taniguchi et al., 2011) crossed with Ai94D (Madisen et al., 2010) and CAG-stop-tTA2 (Miyamichi et al., 2011). We then implanted a glass cylinder, closed at the inner end with a glass coverslip, on top of the LGN (Figure 3A). This preparation allowed visualization of GCaMP6s-expressing neurons in a field of view of 450 x 450 µm (Figure 3B). We recorded visually evoked responses from dLGN interneurons (Figure 3C) to a set of stimuli consisting of black-and-white gratings drifting in 8 different directions at 3 different velocities (400, 1200, 2400 µm/s on the retina).

**Figure 3:**
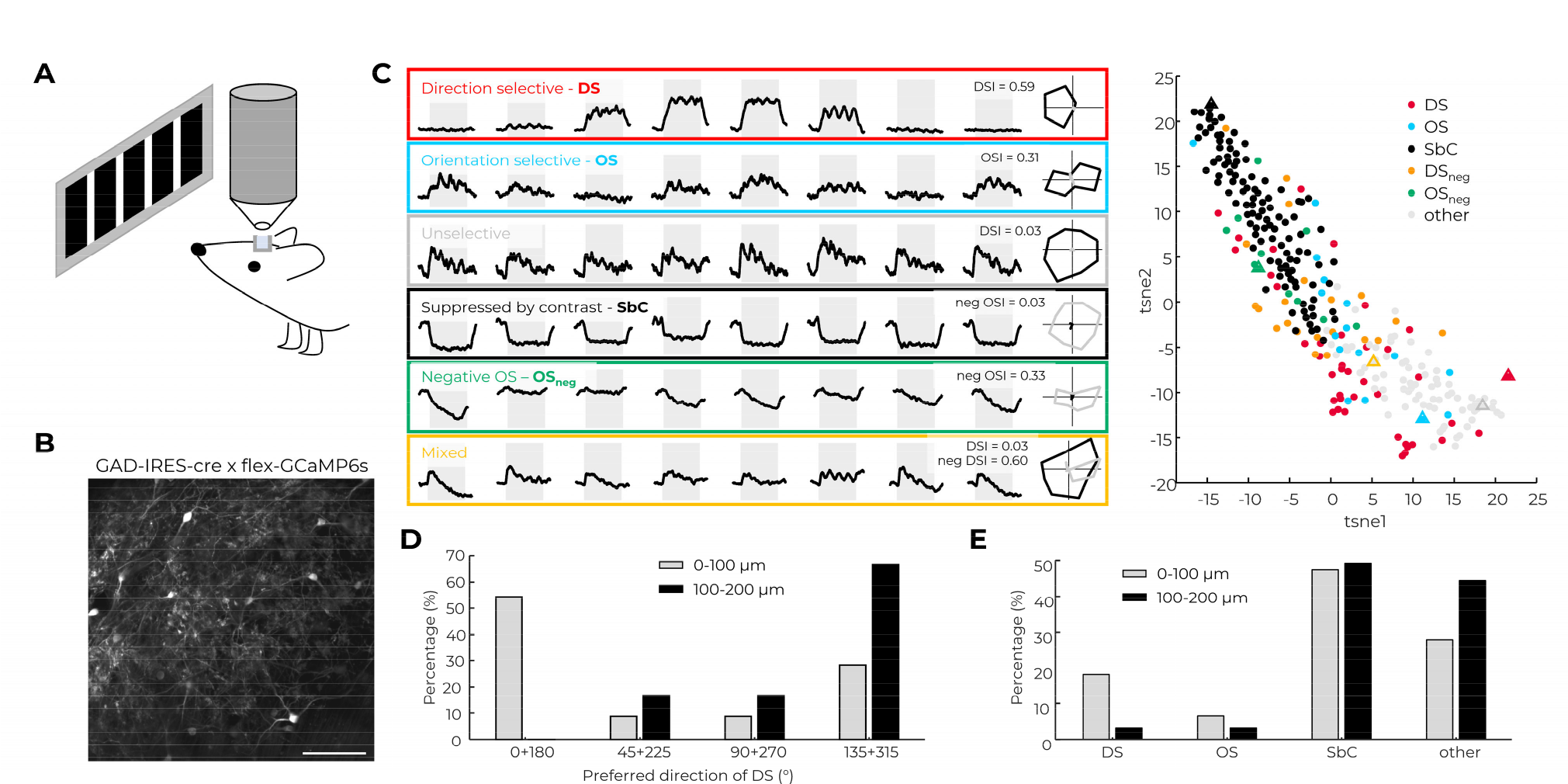
In vivo, dLGN interneurons are specialized towards diverse visual features. **A:** In vivo two-photon calcium imaging in the dLGN is performed via a water-filled glass cylinder implanted on top of the left dLGN. The mouse is head-fixed and receives visual stimulation on an LED screen. **B:** Example field of view with GCaMP6s-expressing dLGN interneurons. Scalebar: 100 µm. **C:** Responses of 6 example interneurons (left panel) to gratings drifting in 8 different directions with 400 µm/s during the shaded time interval. Each polar plot (right of left panel) indicates the positive (black) and negative (gray) response amplitudes for the respective motion direction. The 48-dimensional response vectors (positive and negative amplitudes to 8 directions with 3 velocities) for 278 interneurons is shown in a 2D t-SNE plot (right panel). Triangles indicate the response vectors of the 6 example interneurons shown left in matching colors. **D:** Histogram of the preferred directions of all direction-selective (DS) interneurons (DSI>0.3) at two different imaging depths. **E:** Histogram of response categories: direction selective (DS, DSI>0.3), orientation selective (OS, OSI>0.3), suppressed-by-contrast (SbC, >50% average responses suppressed), and all others, at two different imaging depths. **C,D,E:** Interneurons recorded in 8 wild-type mice.

We observed a wide range of different dLGN interneuron visual responses to the 24 different stimuli. Some interneurons were unselective for the presented stimuli, others were direction selective, orientation selective, or suppressed-by-contrast (Figure 3C). Yet other interneurons had mixed responses such as positive-and-negative or phasic-and-tonic responses to the different stimuli, which could result in suppressed-by-contrast responses with orientation or direction selectivity of the negative amplitudes (Figure 3C). We quantified orientation or direction selectivity of negative amplitudes analogously to positive amplitudes and refer to the selectivity indices as positive or negative direction selectivity index (DSI, DSI_neg_) or positive or negative orientation selectivity index (OSI, OSI_neg_). We then performed dimensionality reduction of the 48- dimensional response vectors defined by the positive and negative response amplitudes to the 24 stimuli (example in Supplemental Figure 3). We found no clear separation of the response vectors into clusters but rather they formed a continuum distributed across a wide feature space (Figure 3C, Supplemental Figure 3). Likewise, the selectivity indices to direction or orientation were widely distributed without clear separation into clusters. The suppressed-by-contrast feature exhibited a bimodal distribution, but the two distributions overlapped, and interneuron examples could be found of intermediate percentages of suppressed response. 15.4% and 19.4% of suppressed-by-contrast interneurons had strong selectivity for direction respectively orientation (Supplemental Figure 3). Altogether, the functional properties of interneurons *in vivo* were diverse, with different features merging into each other.

Previous studies have indicated a dorsoventral gradient from superficial to deeper dLGN for direction-selective retinal ganglion cell projections (Kay et al., 2011; Kim et al., 2008; Krahe et al., 2011; Martersteck et al., 2017; Okigawa et al., 2021; Rivlin-Etzion et al., 2011), as well as an overrepresentation of direction/orientation-selective responses in the dorsolateral dLGN (Piscopo et al., 2013; Román Rosón et al., 2019), in particular an overrepresentation of horizontal motion in the superficial dLGN (Marshel et al., 2012). Therefore, we asked whether dLGN interneuron specialization is also dependent on depth. Superficial interneurons (0-100 µm depth) displayed an overrepresentation of horizontal motion (54.4% preferring 0° or 180° ± 22.5°, Figure 3D). Deeper interneurons (100-200 µm depth) on the other hand showed a significant shift (Fisher’s exact test, p=0.023) away from horizontal motion (0% preferring 0° or 180° ± 22.5°, Figure 3D). The distribution of the four categories − direction selective, orientation selective, suppressed-by-contrast, and others − also changed for deeper relative to superficial interneurons, with a significant drop in the proportion of direction-selective neurons (from 18.1% at 0-100 µm to 3.2% at 100-200 µm, Fisher’s exact test, p=0.002, Figure 3E).

To understand how the functional specialization of individual interneurons is related to the ganglion cell types providing their retinal inputs, we first compared the distribution of the percentage of direction-selective inputs to individual interneurons, as revealed by monosynaptic rabies tracing, to the distribution of the direction selectivity indices of individual interneurons, determined using functional imaging. 18.1% of interneurons in the superficial 100 µm of the dLGN were strongly direction selective (DSI>0.3) in anesthetized animals, whereas 30.1% of superficial interneurons were strongly direction selective in awake animals (Kolmogorov-Smirnov test, p=0.015). Remarkably, the distribution of direction-selectivity indices closely matched the distribution of the percentage of direction-selective inputs received by individual interneurons (Figure 4A). This structure-function correlation indicates that somatic responses reflect the overall distribution of dendritic inputs.

**Figure 4:**
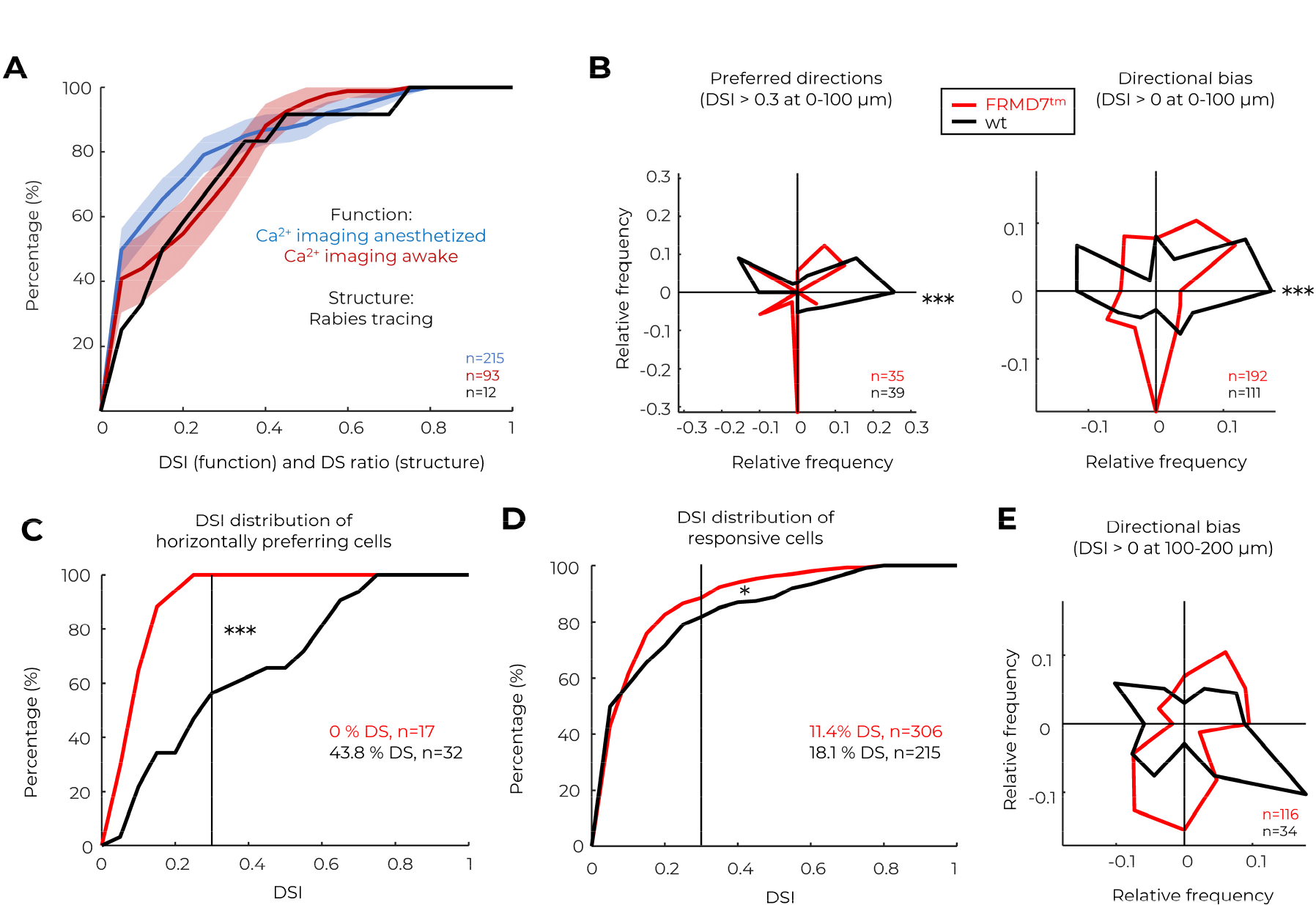
Horizontal direction selectivity of dLGN interneurons is mostly inherited from the retina. **A:** Cumulative histograms of the ratio of direction-selective cells (DS ratio) in individual presynaptic retinal clusters (black, from rabies tracing) and DSI values from calcium imaging in awake (red) and anesthetized (blue) animals. *n*=146 interneurons for calcium imaging in anesthetized, *n*=71 for imaging in awake animals. *n*=638 ganglion cells presynaptic to *n*=12 interneurons for rabies tracing. **B:** Left: Polar plot of preferred directions of direction-selective interneurons (DSI>0.3) in wild-type (wt, black) and FRMD7^tm^ (red) mice. Right: Polar plot of directional bias (direction of summed population vector) of interneurons with DSI>0 in wild-type (wt, black) and FRMD7^tm^ (red) mice. ***: p<0.001, Fisher’s exact test, horizontal directions against all others. **C:** Cumulative histogram of DSI for all horizontal axis-preferring interneurons (population vector direction 0 or 180 ± 15°) in wild-type (black) and FRMD7^tm^ (red) mice. ***: p<0.001, Kolmogorov-Smirnov test. **D:** Cumulative histogram of DSI in wild-type (black) and FRMD7^tm^ (red) mice. *: p=0.031, Kolmogorov-Smirnov test. **E:** Polar plot of directional bias (direction of summed population vector) of interneurons with DSI>0 in wild-type (wt, black) and FRMD7^tm^ (red) mice. **A-E :** Interneurons recorded at 0-100 µm depth (**A-D**) or 100-200 µm depth (**E**), in 8 wild-type and 5 FRMD^tm^ mice.

### Anatomical and functional specializations of dLGN interneurons are causally related

To determine whether the relationship between ganglion-cell input specialization and interneuron functional specialization is causal, we used FRMD7 mutant mice (FRMD7^tm^ mice) in which horizontal direction selectivity is absent in the retina but vertical direction selectivity remains intact (Yonehara et al., 2016). Direction-selective interneurons (DSI>0.3) of FRMD7^tm^ mice showed a significant change in the distribution of preferred directions compared to wild-type mice (Chi-squared test, p<0.001 Figure 4B, Supplemental Figure 4), with a strong reduction in the percentage of neurons preferring horizontal (0 ± 15° and 180 ± 15°) motion directions (from 35.9% to 0% at 0-100 µm depth, Fisher’s exact test, p<0.001). This was not only due to a reduced number of highly horizontally selective interneurons, but the directional bias of less well-tuned (DSI>0) interneurons was also reduced in the horizontal direction (Figure 4B, from 28.8% to 8.9%, Fisher’s exact test, p<0.001), and all interneurons preferring to any degree horizontal motion were less sharply tuned to horizontal motion (Kolmogorov-Smirnov test, p<0.001, Figure 4C, Supplemental Figure 4). This resulted in an overall decrease in the percentage of direction-selective neurons (from 18.1% in wild-type to 11.4% in FRMD7^tm^ mice, Kolmogorov-Smirnov test, p=0.031, Figure 4D, Supplemental Figure 4). Also, deeper interneurons (at 100-200 µm) in FRMD7^tm^ mice had less bias along the horizontal axis and more bias along the vertical axis compared to wild-type mice (Figure 4E). Interestingly, while we did not observe strictly horizontal directions (0° or 180 ± 15°) preferring interneurons, few interneurons strongly selective for less strictly horizontal directions (± 22.5°) remained in FRMD7^tm^ mice, indicating that while most horizontal direction selectivity is inherited from the retina, some cells can recompute it at the level of dLGN. Taken together, the experiments with FRMD7^tm^ mice suggest, at least for direction selectivity, that ganglion-cell input specialization and interneuron functional specialization are causally linked.

### dLGN interneurons display visual features globally

Considering what the function of the strong specializations we observed at interneuron somata might be, it is an intriguing possibility that they may influence local feedforward inhibition at the dendro-dendritic output synapses. A necessary requirement for this would be that the somatic specializations are also present in the dendrites. *In vitro*, it has indeed been shown that dLGN interneurons exhibit backpropagating dendritic spikes (Casale and McCormick, 2011). To test whether *in vivo* the somatic specializations also extend into the dendrites, we annotated the dendrites that ran parallel to the imaging plane for individual dLGN interneurons (Figure 5A). We annotated 1342 dendritic compartments from 227 interneurons of which 1136 fulfilled the signal-to-noise ratio >2.5 criterion. Somatic and dendritic responses were similar (Figure 5A), as reflected by higher correlations within than between interneurons (Figure 5B). Correlations dropped slightly from 0.94 to 0.86 between 1 and 75 µm from the soma (Spearman r^2^<0.04, Supplemental Figure 5), but did not significantly decrease further along the dendrites beyond 75 µm distance (Spearman r^2^<0.005, p>0.3). Average correlations of the response vectors remained higher than 0.8 along the dendrites, both in anesthetized and awake animals (Figure 5C). The percentage of suppressed-by-contrast responses was stable with average deviations less than 5.4% and average absolute deviations less than 15% (Figure 5D, Supplemental Figure 5). The direction- and orientation-selectivity indices of neurons with DSI>0.3 or OSI>0.3 decreased slightly within the first 20 µm but then stabilized along the dendrites (Figure 5E, Supplemental Figure 5). The preferred directions/orientations of the responses were stable with average absolute deviations of less than 25° along the dendrites (Figure 5F, Supplemental Figure 5). Therefore, dLGN interneurons are not only functionally specialized at their soma but also at their dendrites and the two specializations match.

**Figure 5:**
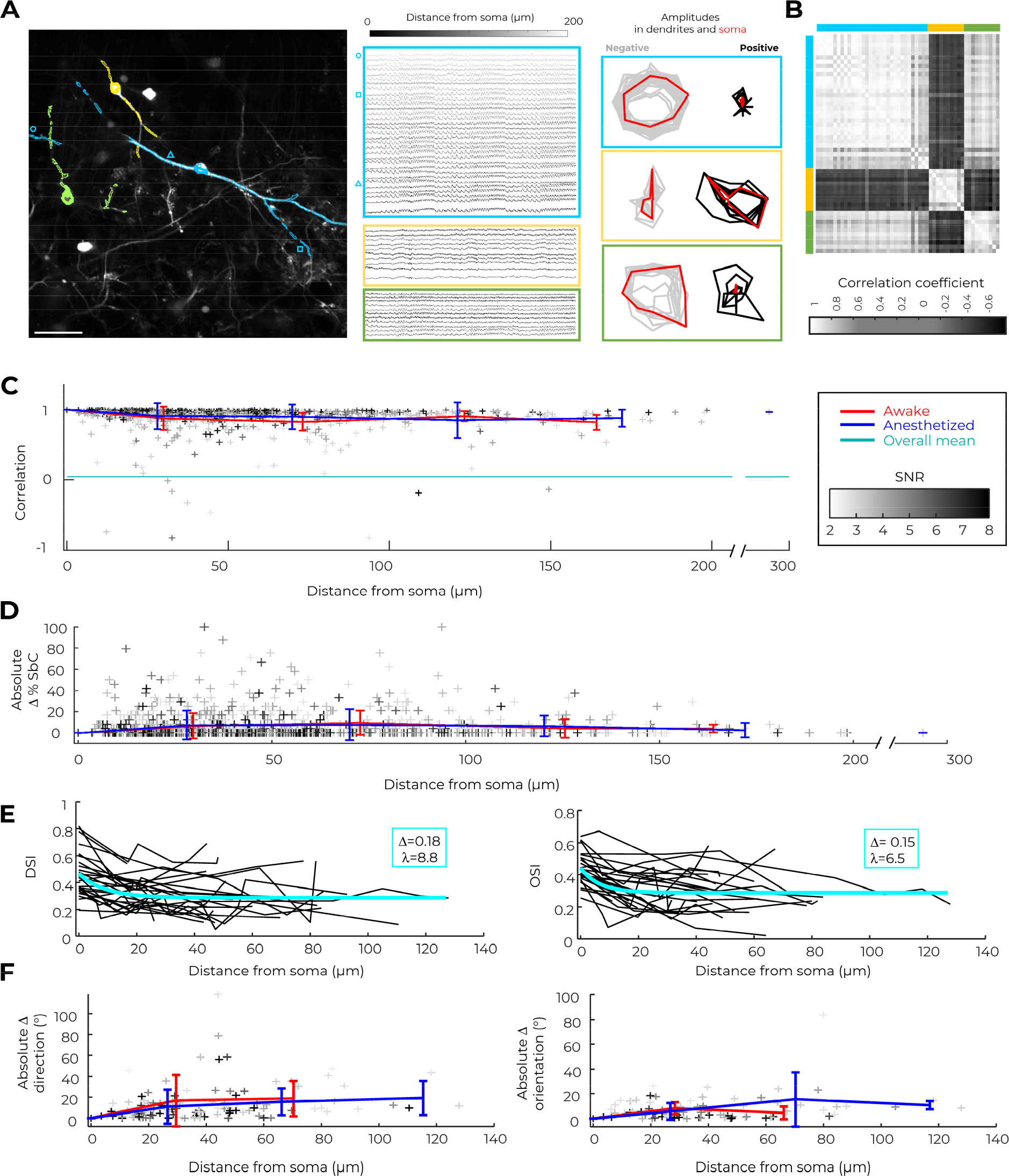
Somatic visual features extend into the dendrites. **A:** Left: three example interneurons with their annotated dendritic regions of interests, recorded in an awake mouse. Scalebar: 50 µm. Middle: calcium responses to visual stimulation recorded at different dendritic positions in each of the three example interneurons. Frame color matches the interneuron color on the left, gray scale indicates Euclidean distance from soma (black = 0 µm). Symbols indicate selected positions labeled on the left. Right: polar plot of negative (left, gray) and positive (right, black) visual response amplitudes to gratings drifting at 400 µm/s velocity in 8 different directions. Each gray/black polar plot corresponds to responses at a different position along the dendrites. Somatic response polar plots are shown in red. **B:** Correlation matrix for the 16-dimensional vectors of positive and negative response amplitudes of the three example interneurons in response to gratings drifting at 400 µm/s velocity in 8 different directions. Colors on the side indicate the identity of the interneurons from A. **C:** Response correlation of all annotated dendritic compartments with their respective soma (using the 16-dimensional response vector), plotted against Euclidean distance from the soma. Individual data points are shaded according to their signal-to-noise ratio (SNR). Red/blue: mean ± standard deviation of data recorded under awake (red) or anesthetized (blue) conditions in 50µm bins. Green line indicates the mean of the full 1136-dimensional correlation matrix of all compartments. For each interneuron, the speed which evoked the largest absolute response was selected. **D:** Absolute differences in the percentage of suppressed-by-contrast (SbC) responses recorded in dendritic compartments with respect to their soma, plotted against Euclidean distance from the soma. Individual data points are shaded according to their signal-to-noise ratio. Red/blue: mean ± standard deviation of data recorded under awake (red) or anesthetized (blue) conditions in 50µm bins. **E:** DSI (left) and OSI (right) values of interneuron dendritic compartments, plotted against Euclidean distance from the soma. Cyan: mono-exponential fit with length constant λ and amplitude Δ as indicated (y = Δ·exp(-x/λ) + y_0_). **F**: Absolute differences in preferred direction (left) or preferred orientation (right) between dendritic compartments and soma, plotted against Euclidean distance from the soma. **E,F:** Included are cells with any compartment showing DSI>0.3 (left) or OSI>0.3 (right). For each interneuron, the speed which evoked the highest DSI (left) or OSI (right) at the soma was selected; see Supplemental Figure 5 for all speeds. Red/blue: mean ± standard deviation of data recorded under awake (red) or anesthetized (blue) conditions in 50µm bins. **C-F:** Interneurons recorded in 5 wild-type, 5 hemizygous FRMD7^tm^ und 3 heterozygous FRMD7^tm^ mice.

## DISCUSSION

We performed single-cell-initiated rabies tracing from and two-photon imaging of dLGN inhibitory interneurons *in vivo* in order to understand how they integrate retinal inputs, both anatomically and functionally. We found that individual dLGN interneurons are both anatomically and functionally specialized to encode diverse visual features, forming a continuum of responses to the same stimuli.

### ‘Multiplexor’ versus ‘Selector’ interneurons

Due to the triadic output arrangements and predicted electrotonic attenuation of their dendrites, interneurons in the dLGN have been thought to act as multiplexors (Bloomfield and Sherman, 1989; Cox et al., 1998; Crandall and Cox, 2013). In such a ‘Multiplexor’ model (Figure 6A), the dendritic triads across the dendritic tree of interneurons act as independent units, receiving input from and shaping the thalamocortical outputs belonging to different retinal channels. In this model, dendrites of ‘Multiplexor’ interneurons would sample a multitude and presumably a random selection of retinal input channels. However, our anatomical and functional results are most consistent with dLGN interneurons being ‘Selectors’. In a ’Selector’ model (Figure 6B), the interneurons receive inputs from a defined subset of retinal channels, which causes the interneuron to respond selectively to the visual feature defined by the given subset of retinal channels. This visual feature arising in the soma of a ‘Selector’ interneuron may back-propagate into the interneuron dendrites and result in a cell-wide functional specialization. The ‘Selector’ model is supported by the following findings. First, the number of ganglion cell types providing input to most dLGN interneurons was smaller than expected by a random draw. Retinal inputs to interneurons displayed a similar degree of specialization as retinal inputs to thalamocortical neurons, reflected by the distribution of specialization scores for ganglion cell types (Figure 2D), cell-type dominances (Figure 2E), and preference indices for transient versus sustained inputs (Figure 2H). As an example, we found interneurons that were highly specialized for receiving input from direction-selective ganglion cells (Figure 2F,I). Second, dLGN interneurons displayed a wide variety of response selectivities *in vivo*, including highly specialized responses such as direction or orientation selective responses. The distribution of direction-selectivity indices of dLGN interneurons *in vivo* matched the distribution of inputs from direction-selective ganglion cells to individual dLGN interneurons (Figure 4A). Third, horizontal direction selectivity of LGN interneurons was strongly reduced in mice lacking horizontal direction selectivity in the retina (Figure 4B,C,E), confirming a causal relationship between presynaptic retinal specialization and the postsynaptic visual feature. Fourth, the selectivities of dendrites of individual dLGN interneurons and their somata were similar (Figure 5), consistent with a cell-wide functional specialization.

**Figure 6:**
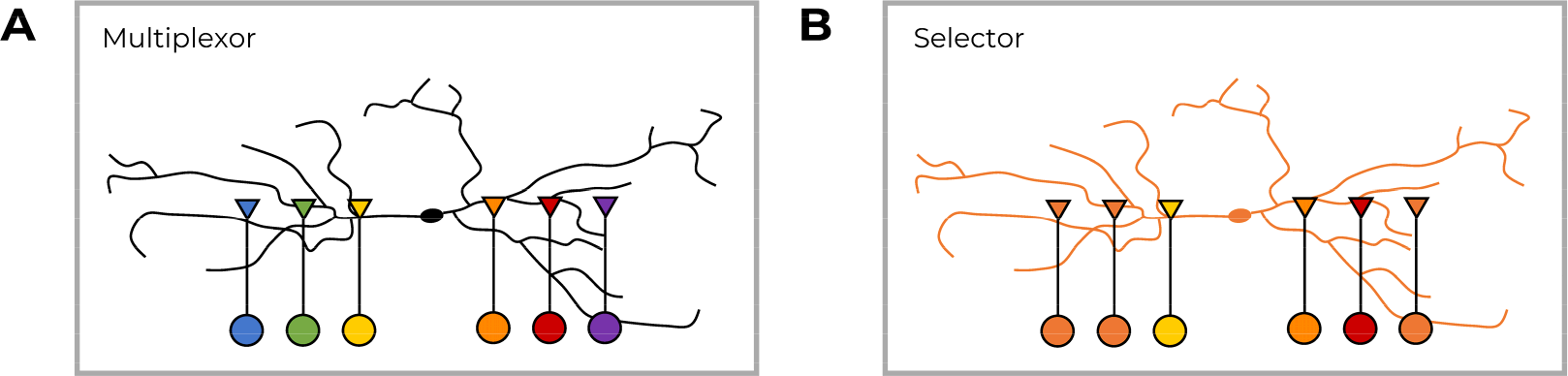
Functional models of dLGN interneurons. **A:** Multiplexor model, with many independent and functionally distinct units along the dendrites. Triangles symbolize the triadic units consisting of retinal ganglion cell-inputs to both the interneuron dendrite and the thalamocortical neuron (not shown), which in turn receives dendro- dendritic inhibitory synapses from the interneuron. **B:** Selector model, in which the interneuron receives functionally specialized inputs from the retina, which dictates its cell-wide functional specialization.

Selection of retinal input channels can result in a visual feature in LGN interneurons that is similar to the dominant retinal input channels but can also result in the computation of a novel visual feature. For example, horizontal direction selectivity is mostly inherited from the retina (Figures 4), but we still observed horizontal direction selectivity in a few interneurons of mice lacking horizontal retinal direction selectivity. This is likely due to *de novo* computation of this visual feature. Additional evidence of *de novo* computation comes from the combined positive/negative and tonic/phasic responses in individual interneurons (Figure 3C), which also suggest that novel features can arise from the combination of distinct retinal channels. Such *de novo* computations may reflect computations based on retinal inputs. Horizontal direction selectivity, for example, could arise from untuned inputs biased towards one direction together with uniformly suppressed-by-contrast inputs that cancel all but one direction (iceberg effect). Alternatively, *de novo* computation may require extraretinal inputs, such as lateral inhibition between dLGN interneurons or inputs from other brain areas such as cortex. Indeed, local inhibitory inputs contributed 7.8% and other extraretinal inputs contributed up to 6.0% of synapses in an dLGN interneuron reconstructed electron-microscopically (Morgan and Lichtman, 2020). These extraretinal inputs could impact the visual response properties of interneurons.

### A possible role for dLGN interneuron specialization: feature-selective attention

What function could ‘Selector’ interneurons fulfill in the dLGN? Extraretinal inputs, such as from cortex, could modulate – activate or inhibit – individual interneurons encoding specific visual features. Since dLGN interneurons have extensive dendritic arbors, in which they display back- propagating action potentials (Acuna-Goycolea et al., 2008; Casale and McCormick, 2011), changes in interneuron activity are globally distributed via their dendrites across the LGN. Importantly, the back-propagating spikes evoke widespread GABA release from interneuron dendrites and inhibitory currents in the postsynaptic thalamocortical neuron dendrites (Acuna-Goycolea et al., 2008). Together with their feature selectivity, these properties could enable dLGN interneurons to perform feature-selective inhibition at the first central stage of vision. Extraretinal inputs activating or inhibiting this feature-selective inhibition could then mediate feature-selective attention, which is thought to gate visual information flow as early as in the dLGN (Ling et al., 2015).

The triadic circuit would be beneficial under this hypothesis for the following reasons. Within a triadic motif, a thalamocortical neuron dendrite receives input from the same retinal ganglion cell as the interneuron dendrite, which provides inhibition to the thalamocortical neuron dendrite. Therefore, most inhibitory output synapses will be provided to thalamocortical neurons that receive at least partially overlapping retinal input channels. Furthermore, since thalamocortical neurons are electrotonically compact and inhibitory inputs by dLGN interneurons arrive at their proximal dendrites, feature-selective inhibition will not only act on dendritic triads with matching retinal input to the thalamocortical neuron but will also influence the neuron’s somatic response and, therefore, its output towards the cortex. These considerations lead to two scenarios.

In the first scenario, the ‘Selector’ interneuron inhibits a thalamocortical neuron that receives the same retinal input channels. This set of channels can be dominated by the same retinal channel or driven by a combination of channels, resulting in a similar mixed feature in both the interneuron and the thalamocortical neuron. Inhibition/activation of this individual dLGN interneuron by extraretinal inputs would then result in disinhibition/suppression of the thalamocortical neuron.

In the second scenario, the ‘Selector’ interneuron inhibits a thalamocortical neuron that receives a partially overlapping but different combination of retinal inputs channels. In this case, the interneuron activity would modify the visual feature encoded by the thalamocortical neuron by activating/suppressing a subset of its combined inputs. Depending on the constellation, this can activate/suppress the feature encoded by the interneuron, while leaving other features encoded by the thalamocortical neuron intact. For example, if a direction-selective interneuron contacts a thalamocortical neuron displaying direction selectivity to the same direction as the interneuron plus orientation selectivity to an oblique axis, the inhibition would affect only the first part of the feature while leaving the second intact.

Likewise, a thalamocortical neuron receiving retinal input channels that do not overlap with the channels received by the interneuron would not receive the inhibition by this interneuron in the first place and consequently would not be influenced. Altogether, the triadic arrangement of synapses increases the likelihood that an interneuron inhibits thalamocortical neurons encoding a similar or the same feature, while reducing the probability of inhibiting thalamocortical neurons encoding distinct features. Such a circuit mechanism ensuring similarity of information content could support the gating of individual visual features by dLGN interneurons, which might be necessary for processes like feature-selective attention.

In addition to dendritic outputs, feature-selective inhibition could also be distributed via the less-prevalent axonal or non-triadic dendritic output synapses of the interneurons. Whether these outputs contact dendrites of thalamocortical neurons with different or similar feature selectivity remains to be shown. Feature-selective inhibition could also cause a modification of visual information that contributes to the emergence of new features (see ‘iceberg effect’ above), rather than providing a simple gating function.

Feature-selective activation of the interneuron dendrites could play an additional role. As in many other cell types, feature-selective dendritic spikes could potentially contribute to cellular plasticity, the absence of which in the dLGN was a long-held postulation but has recently been challenged (Jaepel et al., 2017; Sommeijer et al., 2017).

### The cause of cell-wide feature selectivity

The finding that the encoded visual features measured within the dendrites and somata of interneurons display high similarity *in vivo* (Figure 5) is consistent with the existence of back- propagating spikes described *in vitro* (Acuna-Goycolea et al., 2008; Casale and McCormick, 2011). On the other hand, it would also be consistent with the opposite causality, i.e., the somatic responses resulting from the overall distribution of dendritic inputs. These backwards and forwards causal relationships are not mutually exclusive, especially in a ‘Selector’ model with dominant contribution of one retinal channel (respectively a subset of retinal channels with one common visual feature) and consequently high pre- and postsynaptic correlation. Together, postsynaptic excitation and back-propagating potentials will determine the local dendritic excitation and response selectivity, which in turn determines the dendritic inhibitory outputs.

*In vitro*, synchronous activation of multiple convergent retinal ganglion cells is required to evoke active interneuron firing (Acuna-Goycolea et al., 2008). In visual stimulation that covers large areas of the retina, different retinal ganglion cells are activated simultaneously, providing an explanation for the reliable and strong somatic visual responses that we recorded. The heavy attenuation along the interneuron dendrites predicted by computational simulations for subthreshold voltages (Bloomfield and Sherman, 1989), and observed *in vitro* using glutamate uncaging (Crandall and Cox, 2013), likely underestimated the somatic excitation evoked by multiple convergent inputs.

Whether the dendritic signals reflect back-propagation, forward-propagation, or both, dLGN interneurons are functionally specialized on all levels: their retinal inputs (Figure 2), their dendritic excitation (Figure 5), and their somatic responses (Figure 3). Importantly, our data indicate that the specialization towards selected retinal channels causes the somatic (and matching dendritic) visual feature (Figure 4). It remains to be shown to which degree individual triadic elements and distal dendritic compartments could still be electrically isolated from the global dendritic signals in the incoming and/or outgoing direction and could thereby reflect remaining variability from within the subset of selected retinal channels that they receive (e.g., differences between ON-OFF direction-selective, ON direction-selective and JAM-B inputs in Figure 2F). This question will be especially intriguing for interneurons displaying visual feature selectivity resulting from a mix of relatively diverse retinal channels (for example interneurons in Figure 2D with positive z-scores, Figure 2H with low preference index, Figure 3C with negative orientation selectivity and mixed example).

### Transient versus sustained responses

dLGN interneurons display an overall preference for transient retinal inputs (Figure 2G,H, Supplemental Figure 2). Since the majority of their output and retinal input synapses are found in a triadic arrangement (Morgan and Lichtman, 2020), and assuming that the number of contacts an individual retinal axon makes with each postsynaptic neuron is not different between transient and sustained channels, this implies that dLGN interneurons preferentially contact thalamocortical dendrites receiving transient retinal inputs. Interestingly, in the cat A-laminae, f2 terminals from local interneurons have not been found on Y-cells but on the (more sustained) X-cells (Hamos et al., 1985; Sherman and Friedlander, 1988; Wilson et al., 1984). If also in the cat A-laminae, dLGN interneurons prefer transient channels, they could potentially serve to inhibit transient inputs to the otherwise sustained X-cells. Importantly, f2 terminals have been also found on W-cells in the cat (Raczkowski et al., 1988), which might be more comparable to the mouse superficial dLGN, since they also project to superficial layers in cortex (Sherman and Guillery, 2009).

Potentially, the preference for transient features could be specific for the superficial part of the dLGN that we targeted by electroporation. In this respect, it is notable that the thalamocortical neurons from the same area displayed the opposite preference for sustained inputs. Again, this inverted preference pattern might indicate that transient inputs to cells otherwise preferring sustained inputs are inhibited.

Interestingly, binocular dLGN interneurons display an asymmetric preference pattern for retinal input. Transient retinal channels are overrepresented for contralateral inputs (Figure 2G,H), which constitute most of its inputs (Figure 1E). On the other hand, sustained retinal channels are overrepresented for ipsilateral inputs, which form the minority of inputs. This finding is noteworthy in that the preference of ipsilateral inputs for sustained information matches the preference of ipsilateral inputs in thalamocortical neurons. While the functional role of binocular processing in the dLGN is still debated and deserves further experimental study, we argue that the asymmetric arrangement of retinal information in contra and ipsilateral inputs is not consistent with the hypotheses that ipsilateral inputs to contralateral-input-dominant cells are either developmental remnants that escaped pruning, or serve as a silent backup that is un-silenced by activity-dependent mechanisms if the contralateral eye is deprived (Bauer et al., 2021).

### Superficial versus deep dLGN

By *in vivo* calcium imaging, we were able to image up to 300 µm from the surface of the dLGN, thus covering large parts of the dLGN (∼500 µm deep). Since electroporations are visually guided, they are biased towards the more superficial/dorsal part of the dLGN. Even in this subset of the interneuron population, which underestimates the full variability, we found a large variety of retinal input specializations, corroborating the conclusion that interneurons encode diverse features.

Visual features are not represented evenly across the dLGN. Axons of certain direction-selective retinal ganglion cell types ramify more in the superficial/dorsal dLGN (Kay et al., 2011; Kim et al., 2008; Krahe et al., 2011; Martersteck et al., 2017; Rivlin-Etzion et al., 2011), where also tectogeniculate inputs ramify (Bickford et al., 2015; Grubb and Thompson, 2004; Reese, 1988), and superficial direction-selective dLGN neurons show a preference for motion along the horizontal axis (Marshel et al., 2012). If feature-selective inhibition is indeed, as we hypothesize, important for gating individual visual features, the inhibition needs to be provided only to areas of the dLGN in which ganglion cell axon terminals encode features similar to those of the respective interneuron. Consistent with this, direction-selective interneurons and interneurons preferring motion along the horizontal axis were more prevalent in the superficial than in the deep dLGN (Figure 3D,E).

Intriguingly, we found a higher percentage of direction-selective inputs to interneurons than to thalamocortical neurons described previously (Rompani et al., 2017). There are two possible explanations for this. First, it is possible that the previous selection of electroporated thalamocortical neurons in the superficial dLGN (above the ipsilateral projection zone; see Figure S1 of Rompani et al., 2017) was a subset receiving fewer direction-selective inputs than other superficial thalamocortical neurons, while the dLGN interneurons targeted here, in particular due to their large size, are more representative of the entirety of inputs to the superficial part of dLGN. This interpretation would imply that thalamocortical neurons were sampled from a superficial patch that received fewer direction-selective inputs than the remaining “shell” and indicate that the functional subdivisions of the dLGN go beyond a simple shell-core dichotomy. Second, alternatively or additionally, it is possible that direction-selective retinal inputs could indeed have an overall preference for interneurons over thalamocortical neurons. In this respect, it should be noted that the previous evidence for dLGN thalamocortical neurons in the shell being specialized for direction-selective inputs was based on three arguments: First, several direction-selective retinal ganglion cell types preferentially project to the superficial dLGN (Kay et al., 2011; Kim et al., 2008; Krahe et al., 2011; Martersteck et al., 2017; Rivlin-Etzion et al., 2011, but see also Jiang et al., 2022; Kay et al., 2011 for direction-selective retinal ganglion cell types that target deeper parts). These studies indicate that the superficial dLGN receives more direction-selective ganglion cell inputs than the deeper dLGN. However, axonal projection patterns do not distinguish postsynaptic cell identity. Second, rabies tracing from upper-layer V1 projecting, superficial dLGN neurons labeled almost exclusively direction-selective retinal ganglion cells (Cruz-Martín et al., 2014). This rabies tracing utilized G-coated rabies infection in cortex and relied on G-protein delivery to dLGN neurons via AAV infection, which does not rule out multi-synaptic jumps via interneurons. A third piece of evidence comes from a report that direction-selective neurons in the superficial dLGN prefer horizontal motion (Marshel et al., 2012). The percentage of direction-selective neurons in Marshel et al. was 5% of neurons with DSI>0.5 in the superficial dLGN, compared to 11.2% of interneurons with DSI>0.5 (18.1% for DSI>0.3, 28.4% for DSI>0.2, Supplemental Figure 4) amongst responsive neurons in the upper 100 µm in our study. For comparison, direction-selective ganglion cells that project to the LGN make up 17-35% of all retinal ganglion cells (Baden et al., 2013; Kay et al., 2011; Liang et al., 2018; Peng et al., 2017; Sabbah et al., 2017, DSI criteria 0.2-0.3). Marshel et al. used OGB for imaging activity, which does not distinguish between thalamocortical neurons and interneurons. Taken together, both potential causes – a spatial inhomogeneity of direction-selective thalamocortical neurons and/or direction-selective ganglion cell axons preferentially targeting dLGN interneurons – are possible scenarios, and the degree to which they contributed to the surprisingly low number of direction-selective retinal inputs to thalamocortical neurons compared to interneurons remains to be explored.

### Interneuron cell types and functional diversity

In some brain regions, such as the retina, cell types defined morphologically or transcriptomically are functionally also distinct units. In the dLGN, morphological (Gabbott and Bacon, 1995; Montero and Zempel, 1985) and electrophysiological (Dubin and Cleland, 1977; Leist et al., 2016) studies found independent evidence for only two interneuron types, and single-cell transcriptomic studies identified three transcriptomic types (Bakken et al., 2021). In contrast, we found that mouse dLGN interneurons display many more than three response types to moving grating stimuli. Indeed, our data suggest that dLGN interneurons, rather than being separable into a few functional subtypes, represent a functional continuum with each individual interneuron representing a distinct feature (Figure 3C). However, how individual transcriptomically/morphologically/electro- physiologically defined subtypes map onto this functional response space, and whether the functional response space differs between species, remain to be explored.

In the retina, members of each functional cell type tile the visual space. This arrangement provides the retina with translation invariance. Translation invariance is necessary to recognize the same object in different locations of the visual field. Could individual LGN interneuron response types be required to be present in tiles in order to preserve translation invariance? In our functional imaging data, individual dLGN interneurons span a wide range of visual features (Figure 3C). At the same time, individual interneurons cover large parts of the mouse dLGN, at least half of the visual field (Figure 1C; Morgan and Lichtman, 2020). Therefore, dLGN interneurons provide their feature-selective inhibition globally and thus few or even individual interneurons could preserve translation invariance – highlighting how different the dLGN circuits are from that of the retina.

## METHODS

### Animals

Animals were used in accordance with standard ethical guidelines as stated in the European Communities Guidelines on the Care and Use of Laboratory Animals, 86/609/EEC and FSVO Ordinance on Laboratory Animal Husbandry, the Production of Genetically Modified Animals and Methods of Animal Experimentation (Swiss Animal Experimentation Ordinance) SR 455.163. Experiments were approved by the Veterinary Department of the Canton of Basel-Stadt. Animals for electroporation were 7 females and 6 males, 22 to 60 days old (24 to 34 days old at rabies injection for the single-cell initiated tracing, Supplemental Figure 1), from a GAD67-EGFP line (Tamamaki et al., 2003) or a GAD65-IRES-cre line (Taniguchi et al., 2011) crossed to the EYFP-reporter line (Ai3, Madisen et al., 2010). Animals for imaging were 2 to 6 months old: 4 males and 4 females from GAD65-IRES-cre (Taniguchi et al., 2011) crossed to Ai94D (Madisen et al., 2010) and CAG-stop-tTA2 (Miyamichi et al., 2011) and referred to as wild-type mice; males from this cross were additionally crossed to Frmd7^tm1a(KOMP)Wtsi^ (Yonehara et al., 2016; X-linked genotype) hemizygous females to obtain 5 hemizygous males, referred to as FRMD7^tm^. A further 3 heterozygous FRMD7^tm^ females were not included in the wild-type or FRMD7^tm^ mice categories, because they exhibited an intermediate phenotype (Supplementary Figure 4). Animals were maintained on a 12-h light/dark cycle, fed with irradiated food (KLIBA NAFAG irradiated rodent breeding diet 3302.PM.V20, Provimi Kliba AG) *ad libitum* and autoclaved, chlorinated and acidified tap water. Mice were kept in Individually Ventilated Cages (GM 500, Tecniplast) with bedding (Lignocel BK8-15, Rettenmaier & Söhne GmbH & Co KG) and nesting/enrichment material (Zoonlab GmbH). Health monitoring was done according to FELASA Guideline 2014.

### Single-cell-initiated rabies tracing

Mice were anesthetized with FMM (fentanyl 0.05 mg/kg, medetomidine 0.5 mg/kg, midazolam 5.0 mg/kg). Dexamethasone (2mg/kg, Sigma D2915) was injected to prevent inflammation. Coliquifilm (S01XA20, Allergan) was applied to the eyes to prevent dehydration. A craniotomy was made on the left hemisphere of the mouse skull. Part of cortex and hippocampus above the left dLGN were aspirated to produce an opening of 3 mm diameter centered on the dLGN. The opening above the dLGN was rinsed with Ringer’s solution (150 mM NaCl, 2.5 mM KCl, 2 mM CaCl_2_, 1 mM MgCl_2_, 10 mM HEPES in ddH_2_O, pH 7.4, 0.2 mm sterile filtered). Gelfoam gelatin sponges (Pfizer 9031508) were used to absorb blood. The animal was placed under a custom-made two-photon microscope equipped with red/green detection channels (Hamamatsu R3896 PMTs) and a red LED light source (630 nm Red LED Array Light Source, LIU630A, Thorlabs). An electroporation solution was made containing 40 µl of intracellular solution (130 mM K-methanesulphonate, 10 mM HEPES, 7 mM KCl, 2 mM Na_2_-ATP, 2 mM Mg-ATP, 0.05 mM EGTA in ddH_2_O, 310 mOsm, pH 7.2), 1.5 µl each of pAAV-EF1a-DIO-TVA-WPRE-hGHpA (in GAD65-IRES-cre) or pCMMP-TVA800 (in GAD67-EGFP), pAAV-EF1a-CVS11-G-WPRE-hGHpA, and pAAV-EF1a-tdTomato-WPRE-hGHpA (all at a final working concentration of ∼0.1 µg/µl), and 2.5 µl Alexa-594 (1 mM in intracellular solution, A-10438, Thermo Fisher). The solution was filtered through a 0.2 mm Ultrafree-MC GV Centrifugal Filter (UFC30GV0S, Millipore). In 3/13 animals with the GAD65-IRES-cre genotype, pAAV-EF1a-CVS11-G-WPRE-hGHpA was replaced by pAAV-Ef1a-DIO-oG-WPRE (Kim et al., 2016); however, presynaptic ganglion cell numbers were not increased by oG (70 ± 18 cells with oG versus 105 ± 28 cells with CVS11G; mean ± sem), possibly because transsynaptic labeling was already saturated with CVS11-G. The electroporation solution was loaded into a glass needle (Standard Wall Borosilicate Tubing with Filament, BF100-50-10, Sutter Instruments, resistance 10-30 MΩ) and placed onto the headstage of an electroporation device (Axoporator, 800A, Molecular Devices) mounted on a manipulator (MPC 200, Sutter Instruments). The electrode was placed on the surface of the dLGN using local landmarks (Supplementary Figure 1A) and a 4x objective (MPlan N 5x/.1NA, Olympus). Green-fluorescent interneurons in the dLGN and the Alexa-filled pipette were visualized by two-photon imaging at 850 nm through a 40x objective (LUMPlanFl 40x/ 0.8NA Water immersion, Olympus) and electroporated using the settings: voltage = -6-14 V, DC offset = 0, train = 10-1000 ms, frequency = 100 Hz, pulse width 50-500 µs. In 10 animals, only one green-fluorescent neuron was electroporated per animal. Two animals in which additionally targeted cells were not filled or did not survive electroporation were also counted as single-cell initiated. In one animal, 4 GFP-positive cells were electroporated; this animal was included only when estimating the overall distribution of presynaptic ganglion cell types (Figure 2G). After retraction of the pipette, the surgical window was filled with KWIK-CAST (World Precision Instruments) or a 3-mm glass cylinder sealed with a 3-mm coverslip (see “calcium imaging”), was implanted. The mouse received buprenorphine (0.1 mg/kg) for postoperative analgesia and was placed in a heated cage to recover. Up to 7 days after the electroporation, the window/plug was removed (pharmacological treatment as above), a pulled glass needle (Premium Standard Wall Borosilicate, Model G100-4, Warner Instruments) was cut at the tip (1-3 MΩ) and loaded with EnvA-coated SADΔG-rabies. The EnvA-coated SADΔG-rabies virus was produced as described previously (Rompani et al., 2017; Wertz et al., 2015) and expressed either mCherry (Rompani et al., 2017) or tagRFP (generously provided by Karl-Klaus Conzelmann). The viral titer ranged from 10^8^ to 10^12^ plaque-forming units/ml. In order to minimize dilution of the rabies virus, Ringer’s media from the LGN surface was removed. Rabies virus (250 nl – 1.2 µl) was injected within 200 µm of the electroporated cell. To allow virus diffusion within the tissue, the pipette was left in place for >10 min before retraction. Mice were sacrificed 10-11 days after rabies injection.

### Immunohistochemistry

After euthanasia, eyes were harvested and fixed in 4% PFA overnight at room temperature (RT). Retinas were dissected and washed 3x in PBS, transferred to 30% sucrose in PBS (w/v), allowed to sink and then subjected to 3 freeze-thaw cycles. Retinas were again washed 3x in PBS and incubated in heavy blocking solution (10% NDS, 1% BSA, 0.5% Triton X-100, 0.01% sodium azide in PBS) for 1 h at RT. Primary antibodies (goat a-ChAT Millipore AB144P 1:200 and chicken a-RFP Rockland 600-901-379 1:1000 or rabbit a-tRFP Evrogen AB233 1:1000) were prepared in light blocking solution (3% NDS, 1% BSA, 0.5% Triton X-100, 0.01% sodium azide in PBS) and retinas treated for 3-14 days at RT. Retinas were then washed 3x with PBS. Secondary antibodies (donkey a-goat Alexa488 A11055 Life Technologies 1:200 and a-chicken Cy3 Jackson F03-165-155 1:200 or a-rabbit Alexa568 A10042 Life Technologies 1:200) were prepared in light blocking solution and retinas were treated for 1-2 h at RT. After 3x washing in PBS, the retinas were mounted with ProLong Gold (P36934 ThermoFisher), with strips of Parafilm (52858-000 VWR) at the coverslip borders to prevent tissue compression.

### Confocal microscopy

Confocal image stacks were acquired using spinning disc microscopes with a CSU W1 dual camera T2 spinning disk confocal scanning unit (Yokogawa), a homogenizer (Visitron), 63x/1.4 or 40x/1.3 Plan-Apochromat oil objectives (Zeiss), and a MS2000 X,Y stage with a Z-Piezo drive (ASI). The upright system was built on an AxioImager M2 microscope (Zeiss) and equipped with two Edge cameras (PCO). The inverted system was built on an AxioObserver (Zeiss) and equipped with two Prime 95B cameras (Photometrics). Images had a pixel size of ∼0.2 µm in xy, 0.25 µm in z. Appropriate illumination and filters were used for Alexa488 (ChAT) and Cy3/Alexa568 (retinal ganglion cells) and a subset of images acquired on a Zeiss LSM 720. Sub-micrometer z-offsets between the red and green channels were detected and compensated posthoc.

### ChAT-band detection

To generate training data, ChAT-bands were manually annotated as two lines in the yz-projections of a subset of confocal stacks. These data were used to train a U-Net (Ronneberger et al., 2015), implemented in Python using Keras/TensorFlow, in order to detect ChAT-bands in 3D stacks. The output was provided as a 3D probability stack. Subsequent analysis was performed in Matlab (R2019b, Mathworks). From the 3D probability stack, the two ChAT-bands were automatically detected in yz-projections as lines of local maxima, resulting in a 2D (xy) matrix of z-coordinates for each band. Correct assignment of the ChAT-bands was manually verified and curated if necessary.

### Classification of retinal ganglion cells

The z-coordinates of the ChAT-bands were used to artificially flatten the image stacks by assigning each pixel its relative position to the ChAT coordinates in z, set to relative position 0 and 1, and interpolating the stack at z-increments of 0.025. 10 strata were defined at regular intervals of 0.25, with strata 3 and 7 centered around the OFF- respectively ON-ChAT coordinates. For the red channel (of labeled retinal ganglion cells), a maximum z-projection was pseudo-colored with respect to the stratum in which the pixel with maximum brightness was located (Figure 2C,F, Supplemental Figure 2). Additional maximum projections per stratum of both channels aided the classification (for example to evaluate co-fasciculation of dendrites with the ChAT-processes). Based on these visualizations, individual ganglion cells were manually classified by the stratification of their terminal dendrites. Type 12 included cells with stratification in strata 1 and/or 2. If the dendrites also displayed asymmetry characteristic of JAM-B, the cells were classified as 12_asym. Type 89 included the previous types 89_big and 89_PV1 (Rompani et al., 2017), as well as other cells stratifying in strata 8 and/or 9 and/or 10. Type 189 included the previous types 189_big and 189_small, as well as other cells stratifying in 1, 2 and 8 and/or 9 and/or 10. Type 4 included the previous types 4_giPV5, 4_PV5, 4_PVX. Type 37 cells co-stratified with the ChAT- processes in 3 and 7. Type 7 co-stratified with the ON-ChAT stratum 7, with only minor branches to stratum 3 (see examples in Figure 2F). Cells stratifying in 689 were not found amongst the cells presynaptic to dLGN interneurons. Direction selective ratios (Figure 4A) were measured as the ratio of direction-selective presynaptic retinal ganglion cells (defined as co-stratifying with the ChAT-bands or stratifying in strata 1 and 2 and displaying asymmetric dendrites characteristic of JAM-B cells) relative to all classified ganglion cells presynaptic to a single dLGN interneuron. Of *n*=1257 retinal ganglion cells presynaptic to dLGN interneurons, *n*=688 could be classified. This included the *n*=638 classified out of *n*=1157 retinal ganglion cells labeled by *n*=12 single-cell-initiated rabies tracing, plus one experiment with 4 electroporated interneurons. Data for thalamocortical neurons refer to the *n*=245 classified out of *n*=507 total retinal ganglion cells previously reported (Rompani et al., 2017).

### Probabilistic modeling

Monte-Carlo simulations were performed in Matlab (R2019b, Mathworks) to simulate the null hypothesis that presynaptic ganglion cells were randomly drawn from a given distribution of cell types. To obtain conservative estimates of specialization, we assumed that the distribution of cell types projecting to the dLGN is not uniformly random, but reflected in the overall distribution of cell types projecting to thalamocortical neurons (which itself is specialized, see Rompani et al., 2017) and the overall distribution of cell types presynaptic to dLGN interneurons (Figure 2G). To compare specialization between dLGN interneurons and thalamocortical neurons, we estimated the overall distribution of cell types by the empirical distributions p_TCN_ and p_IN_ found for thalamocortical neurons, respectively interneurons, and calculated a weighted average (p_TCN_ · W_TCN_ · N_TCN_ + p_IN_ · W_IN_ · N_IN_), with N_TCN_ = 20.3, N_IN_ = 96.4 being the average empirical numbers of presynaptic retinal ganglion cells per thalamocortical neuron / interneuron and W_TCN_=0.8, W_IN_=0.2 to account for the higher prevalence of thalamocortical neurons. From this distribution, the expected number of cell types present in any given number of presynaptic cells and its standard deviation were simulated. Specialization z-scores were calculated as the difference between empirically found and expected number of cell types divided by the simulated standard deviation; the number of classified ganglion cells was taken as the total number of cells for this calculation. Use of the weighted empirical distribution, which included the specialized distribution of thalamocortical neurons, resulted in conservative estimates for interneuron specialization and 2 positive z-scores >1 (“systematically more random than expected”), which are closer to the random expectation if the simulation is based on the empirical distribution p_IN_ found in interneurons (Supplemental Figure 2).

### *In vivo* two-photon calcium imaging

Surgery was performed (see “Single-cell initiated rabies tracing”, same pharmacological treatment) to obtain a 3-mm cranial window over the left dLGN, with overlying parts of cortex and hippocampus removed. A custom-made cylinder of borosilicate glass (OD 3 mm, ID 2.55 mm, length 1.6 mm), sealed with a 3-mm glass coverslip (CS-3R-0, Warner Instruments), was implanted and fixed to the skull with superglue (Pattex Ultragel). A custom-made aluminium headplate was attached to the skull with superglue and dental cement (Paladur). Two-photon calcium imaging was performed after recovery from the surgery, at the earliest 1 day later. Imaging was performed on 1-7 days up to 30 days post implantation. Mice were head-fixed via the headplate. For anesthetized recordings, mice were injected with 1.25 mg/kg chlorprothixene prior to recording, lightly anesthetized with ∼0.5% isoflurane during recording and kept on a heating pad. For awake recordings, mice were accustomed to and later imaged while being head-fixed and freely running on a wheel. An LED screen (52.5 cm wide, 29.5 cm high, 15 cm distance) was placed in front of the right eye. The glass cylinder was filled with ddH_2_O prewarmed to 38°C. Two-photon imaging was performed using a FemtoSMART resonant-galvo scanning microscope equipped with a 16x Nikon water immersion objective (N16XLWD, 0.8 NA, 3 mm WD), which was warmed and light-shielded. Data were acquired at 920-nm illumination with up to 60 Hz, 0.4-1.8 µm per pixel, up to 475 x 475 µm field of view. For visual stimulation, white-black gratings were shown that moved in 8 different directions with 3 different velocities (400/1200/2400 µm/s on the retina, white bar width 10°, 40° per cycle). A gray screen with the same average (25%) luminance was shown between moving gratings.

### Data analysis

Data was processed by custom-written software in Matlab (R2019b, Mathworks). Bidirectional scanning artifacts were corrected and rigid motion-correction was performed. Data were filtered by a moving average filter of 500 ms length. Regions of interest (ROIs) were automatically detected based on local correlations after filtering, briefly: A matrix of local correlations was calculated. ROIs were initiated at local maxima of the correlation matrix and extended to neighboring pixels if a correlation threshold was exceeded, which depended both on local correlations of the seeding pixel and the local background correlations. Local background was defined as the filtered average signal of the 10 darkest pixels within 15 µm distance. Local background was subtracted for all ROIs (even though it contained visible signals only in a few very densely labeled regions). For each recording (usually 4 repetitions of moving gratings in 8 different directions at 3 different velocities), ROIs corresponding to putative somata were manually selected from an overlay of the 95^th^ percentile projection and the local correlation matrix. For all putative somata, local correlation matrices and maximum projections over the frames with highest activity at this ROI were inspected and discarded if they were not indicative of a soma (Supplemental Figure 6). To avoid duplication of cells, corresponding imaging regions recorded consecutively or on different days were aligned, matching ROIs were determined (Supplemental Figure 6) and the data for each ROI was concatenated, respectively the dataset with the highest signal-to-noise ratio (SNR) was selected. SNR was defined as maximum response amplitude divided by the standard deviation of the baseline. ROIs were excluded if strong trial-to-trial variability unrelated to the stimulus was present or if the SNR did not exceed 2.5. Baseline was defined as median signal within 1.75-0.25 s before stimulus onset. The median response of raw signals for each stimulus was determined after baseline subtraction, mostly from 4 (range 3-15) stimulus repetitions. The median response was filtered, its positive response P_i_ (maximum), negative response N_i_ (-minimum), and average across the stimulation period were calculated. A response was counted as suppressed if the average was below baseline. If >50% of the 24 average responses were negative, the cell was classified as suppressed-by-contrast (SbC). For each velocity, the direction- and orientation-selectivity indices were calculated from the 8-dimensional tuning vectors P_1-8_ and N_1-8_ (after setting negative values to zero) to directions ϑ_1-8_ based on circular variance:

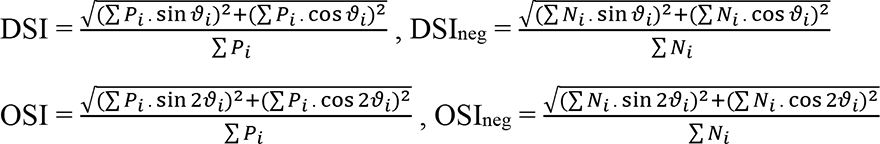

If max(P_i_)<max(N_i_), DSI was set to zero, if max(P_i_)>max(N_i_), DSI_neg_ was set to zero, which systematically underestimates the DSI of negatively responding, respectively the DSI_neg_ of positively responding interneurons. Preferred directions/orientations were defined as the angle of the population vector (complex phase for direction, half the complex phase for orientation) and were corrected for the angle between the mouse eye main axis and screen horizontal axis (Supplemental Figure 6). All polar plots shown display responses as ΔF/F_0_, with F_0_ defined as smallest baseline of the 24 average responses. For speed-specific plots, SNR and percentage suppressed by contrast were calculated per velocity, and only data with SNR>2.5 were included. For calculating speed-independent DSI, respectively OSI per interneuron, the speed which evoked the highest DSI, respectively OSI was selected. For calculating speed-independent correlations of dendritic with somatic responses, the speed which evoked the maximum response (positive or negative) was selected. Interneurons with DSI>0.3 and OSI>0.3 were categorized as direction selective. Interneurons suppressed-by-contrast, but DS, OS, DS_neg_ or OS_neg_ were categorized as DS, OS, DS_neg_ or OS_neg_. Imaging depth refers to depth below the optic tract (cells start at 0 µm), which is ∼50 µm below the dLGN surface.

### Dendritic analysis

For all recordings (including the awake condition and recordings in FRMD7^tm^ animals), the overlay of the 95^th^ percentile projection and the local correlation matrix were inspected for cells with dendrites visible in the same imaging plane. From the automatically detected ROIs, dendrites belonging to a soma were manually annotated. In ambiguous cases, for example to distinguish smaller bifurcations from other crossing dendrites, the overall signals (Figure 5A, middle panel) were inspected for similarity. Only compartments with SNR>2.5 were included in further analysis. The morphology of suppressed-by-contrast interneurons, presumably due to their higher baseline activity, could be reconstructed over the longest distances (up to 300 µm). Distances were measured as Euclidean distance between ROI centers, providing a lower estimate for the path length.

### Statistics

Non-parametric tests (Mann-Whitney-U test, Wilcoxon signed-rank test) were applied for all group comparisons. Fisher’s exact test was applied for 2x2 contingency tables, Chi-squared test for comparing larger contingency tables. For the comparison of continuous distributions, Kolmogorov-Smirnov test was applied. To test whether the distribution of presynaptic cell numbers across the two eyes deviated from a random binomial distribution, the p value was derived from the symmetric confidence intervals of the Monte-Carlo simulated distributions of the given test-statistics (absolute difference between ipsi- and contralateral cell counts). For multiple comparisons (Figure 2D, Supplemental Figure 4), the p-values were Bonferroni-Holm corrected. Pearson correlation coefficients are provided to quantify correlations between response vectors (Figure 5B,C). Spearman rank correlation was used to quantify the non-linear distance-dependence (Figure 5C).

## ACKNOWLEDGMENTS

We thank Markus Rempfler from the Facility for Advanced Imaging and Microscopy (FAIM) of the Friedrich Miescher Institute Basel for implementing the U-Net for machine learning-assisted detection of the ChAT-bands. We thank Santiago Rompani for his advice in performing single-cell-initiated rabies tracing and for sharing material, and Karl-Klaus Conzelmann, Keisuke Yonehara, and Martin Munz also for sharing material. We thank all involved animal caretakers and those who provided technical support at FMI or IOB Basel for their help, especially Basil Thommen, Antonio Martínez Brotons, Nicole Ledergerber, Josephine Jüttner, Brigitte Gross Scherf, Claudia Patino Alvarez, Serena Curtoni, Dimitri Rey, Steven Bourke (FAIM), Enrico Tagliavini, Zoltan Raics and Paul Argast. We thank Helene Schreyer, Katja Kolar and Alex Fratzl for comments on the manuscript. This worked was supported by the following Grants: EMBO (ALTF 519-2016), Marie Skłodowska-Curie actions (707522), Research Fund of the University of Basel (3ZX1414) to FEM; Swiss National Science Foundation Synergia grant (CRSII3_141801), European Research Council advanced grant (RETMUS N°669157, HURET N°883781), Louis-Jeantet Foundation award, Körber Foundation award, Swiss National Science Foundation grant (31003A_182523), and the NCCR ‘Molecular Systems Engineering’ to BR.

## AUTHOR CONTRIBUTIONS

FEM conducted experiments and conceived and conducted data analysis and modeling. BR supervised the work. FEM and BR interpreted the data and wrote the manuscript.

---

**Supplemental Figure 1:**
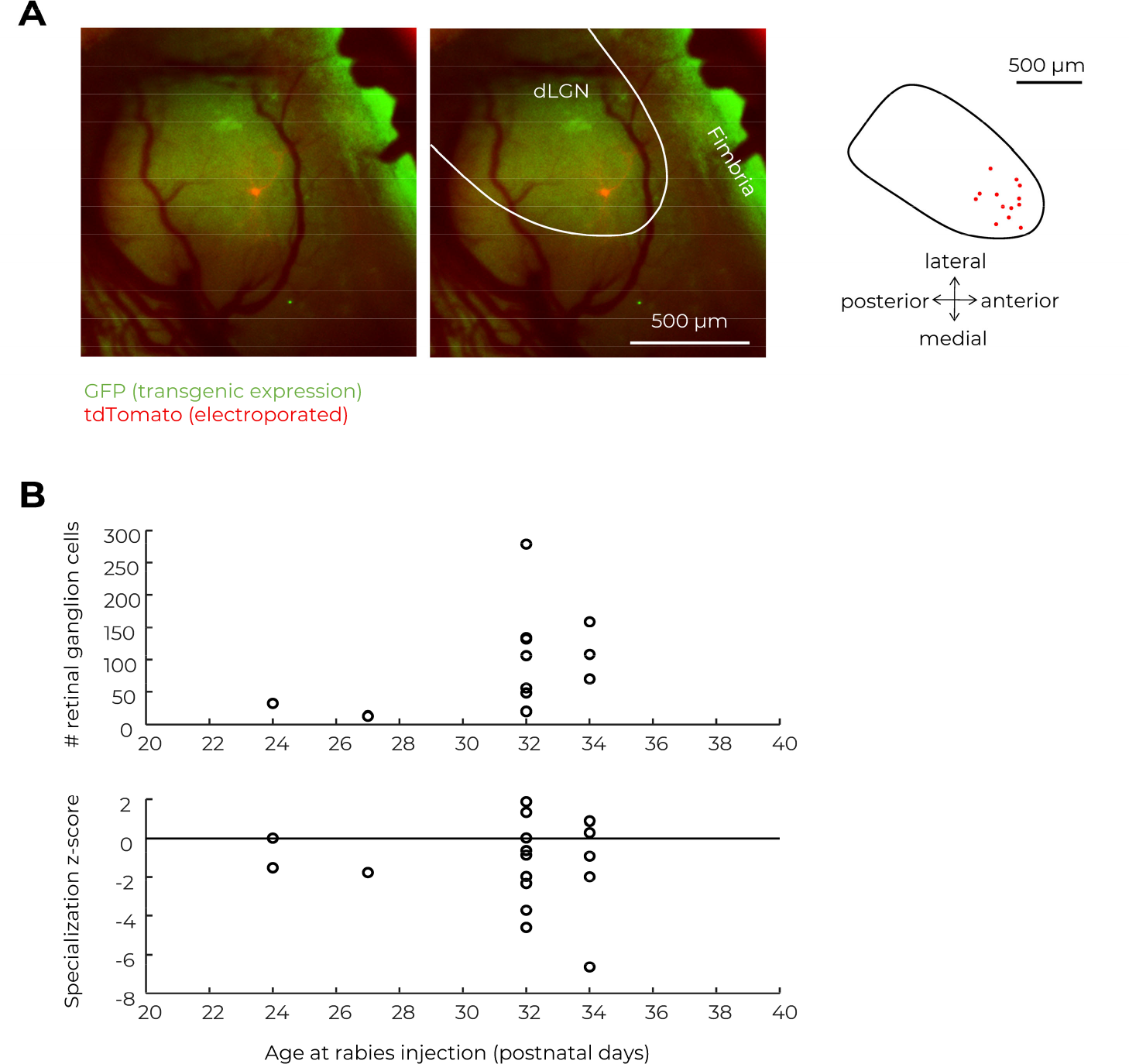
Method details of single-cell-initiated rabies tracing. **A:** To target interneurons in the dLGN by single-cell electroporation, the dLGN was visually identified by the following anatomical landmarks established by intravitreal CTB-injections: (1) 3D-curvature of the thalamic surface and (2) blood vessel pattern provide a putative location, which is confirmed by (3) GFP-expression of genetically targeted GABAergic interneurons which have a higher density in the dLGN compared to surrounding structures. Green: native fluorescence of the GABAergic interneurons. Red: tdTomato expression of the electroporated interneurons at day 3. Left panel shows an epifluorescence image though the implanted window. Middle panel indicates the scaled outlines of the dLGN in white along the border of green expression and the fimbria. Right panel shows the positions of all electroporated interneurons (red) with respect to the dLGN outlines (black). **B:** Measured number of presynaptic retinal ganglion cells (upper panel) and specialization z-scores (lower panel) plotted against the age of mice at rabies injection for all single-cell-initiated rabies tracings.

**Supplemental Figure 2:**
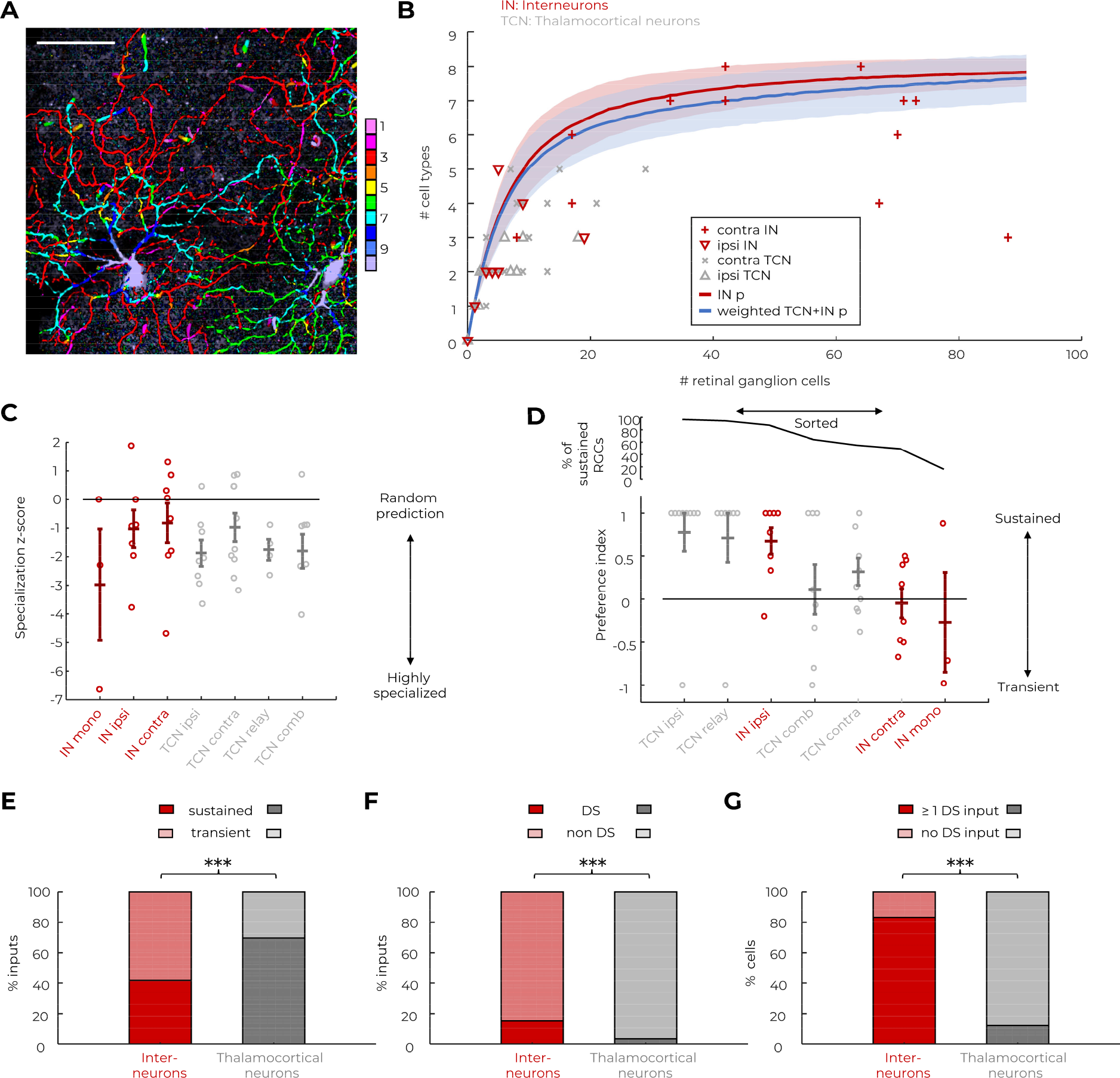
Retinal inputs to dLGN interneurons are specialized. **A:** Two close-by retinal ganglion cells with arbor overlap can be classified as type 37 and type 6 based on the maximum z-projection, in which maxima are pseudo-colored according to their location in strata 1-10 (color bar). Scalebar: 50 µm. **B:** Expected (lines) and measured (symbols) numbers of cell types plotted against the total number of presynaptic cells. Expected numbers were simulated as random draw with replacement (multinomial distribution) based on the weighted average of empirical distributions (’weighted TCN+IN p’, blue) or the empirical distribution for interneurons (’IN p’, red, Figure 2G). Thick line: mean, shaded area: mean ± standard deviation of the simulated numbers. IN: interneurons, TCN: thalamocortical neurons. Symbols indicate the measured numbers of ganglion cell-types in contralateral or ipsilateral retinas presynaptic to the interneurons (’contra IN’, ’ipsi IN’ in red) or thalamocortical neurons (’contra TCN’, ’ipsi TCN’ in gray) from which the retrograde tracing was initiated. **C,D:** Specialization z-scores (**C**), and preference indices (**D**) of individual retinal clusters grouped by the corresponding integration mode. IN: interneurons (red), TCN: thalamocortical neurons (gray). Mono: monocular, ipsi: ipsilateral, contra: contralateral, relay: monocular relay-mode, comb: monocular combination-mode. Relay- and combination-mode refer to the definition in Rompani et al., 2017. Horizontal lines indicate group averages, error bars indicate ± sem. Data groups in **D** are sorted along the x-axis by decreasing percentage of sustained RGCs found in the pooled data per group (upper panel). **E:** Percentages of sustained and transient inputs to interneurons (red) and thalamocortical neurons (gray, Rompani et al., 2017). ***: p<0.001, Fisher’s exact test. **F:** Percentages of direction-selective (DS) inputs to interneurons (red) and thalamocortical neurons (gray, Rompani et al., 2017). ***: p<0.001, Fisher’s exact test. **G:** Proportion of interneurons (red) and thalamocortical neurons (gray, Rompani et al., 2017) receiving at least 1 direction-selective (DS) input. ***: p<0.001, Fisher’s exact test.

**Supplemental Figure 3:**
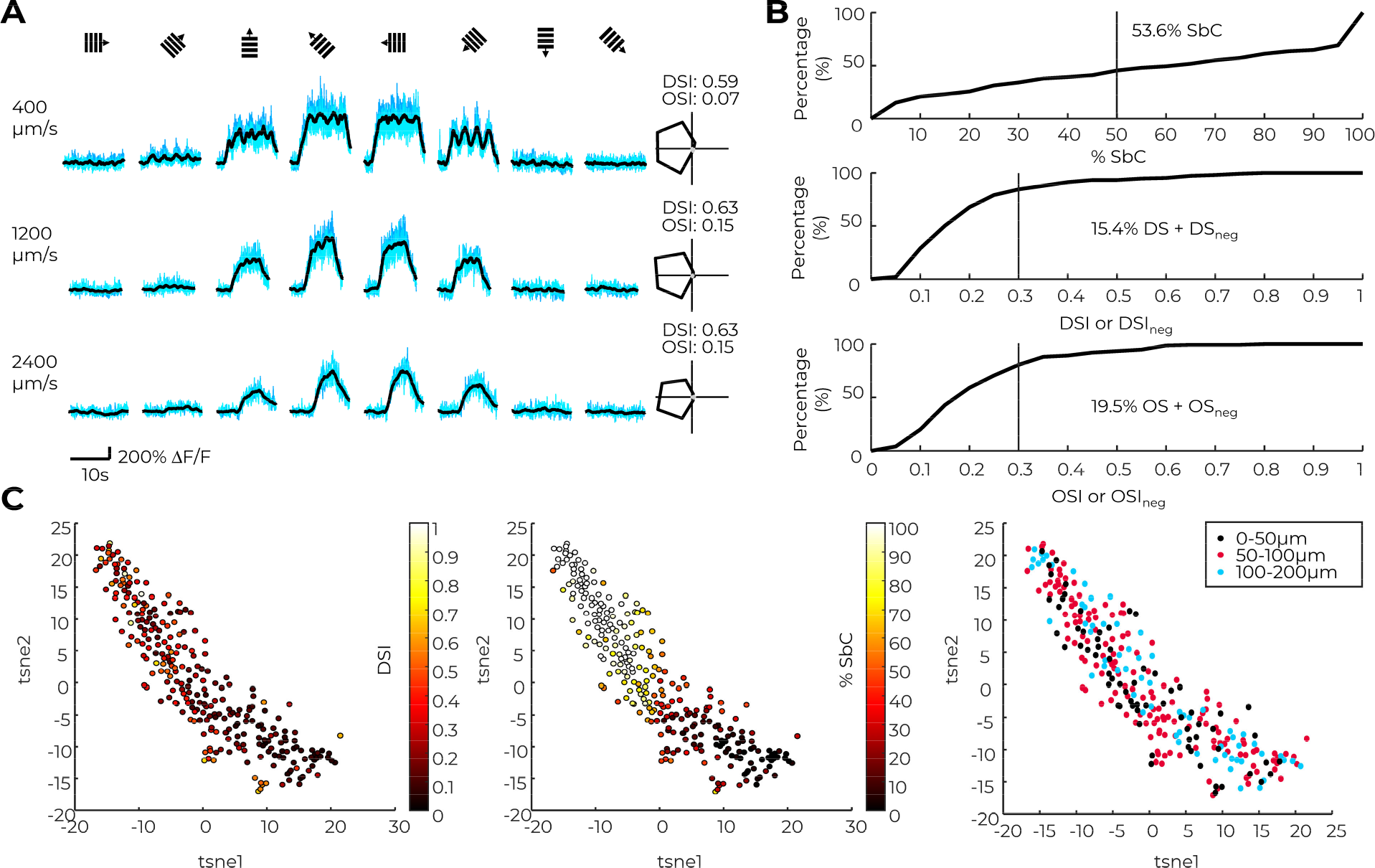
dLGN interneurons display a wide range of visual features. **A:** Example visual responses to the 24 stimuli (gratings drifting in 8 different directions with 3 different velocities). Blue: individual responses. Black: filtered median response. Top to bottom: 400, 1200, 2400 µm/s on the retina. Polar plots on the right display positive (black) and negative (gray, close to zero) response amplitudes plotted with respect to stimulus direction. **B:** Cumulative histograms of suppressed-by-contrast response features. Upper panel: Cumulative histogram of the percentage of responses suppressed by contrast. Middle/lower panel: Cumulative histograms of the direction selectivity indices (middle, DSI and DSI_neg_) and orientation selectivity indices (lower, OSI and OSI_neg_) for interneurons with >50% suppressed-by-contrast responses. **C:** t-SNE plots of the 48-dimensional response vectors (24 stimuli positive and negative responses) pseudo-colored for DSI (left), percentage of suppressed-by-contrast responses (center) and depth from surface (right). **B-C**: Interneurons recorded in 8 wild-type mice.

**Supplemental Figure 4:**
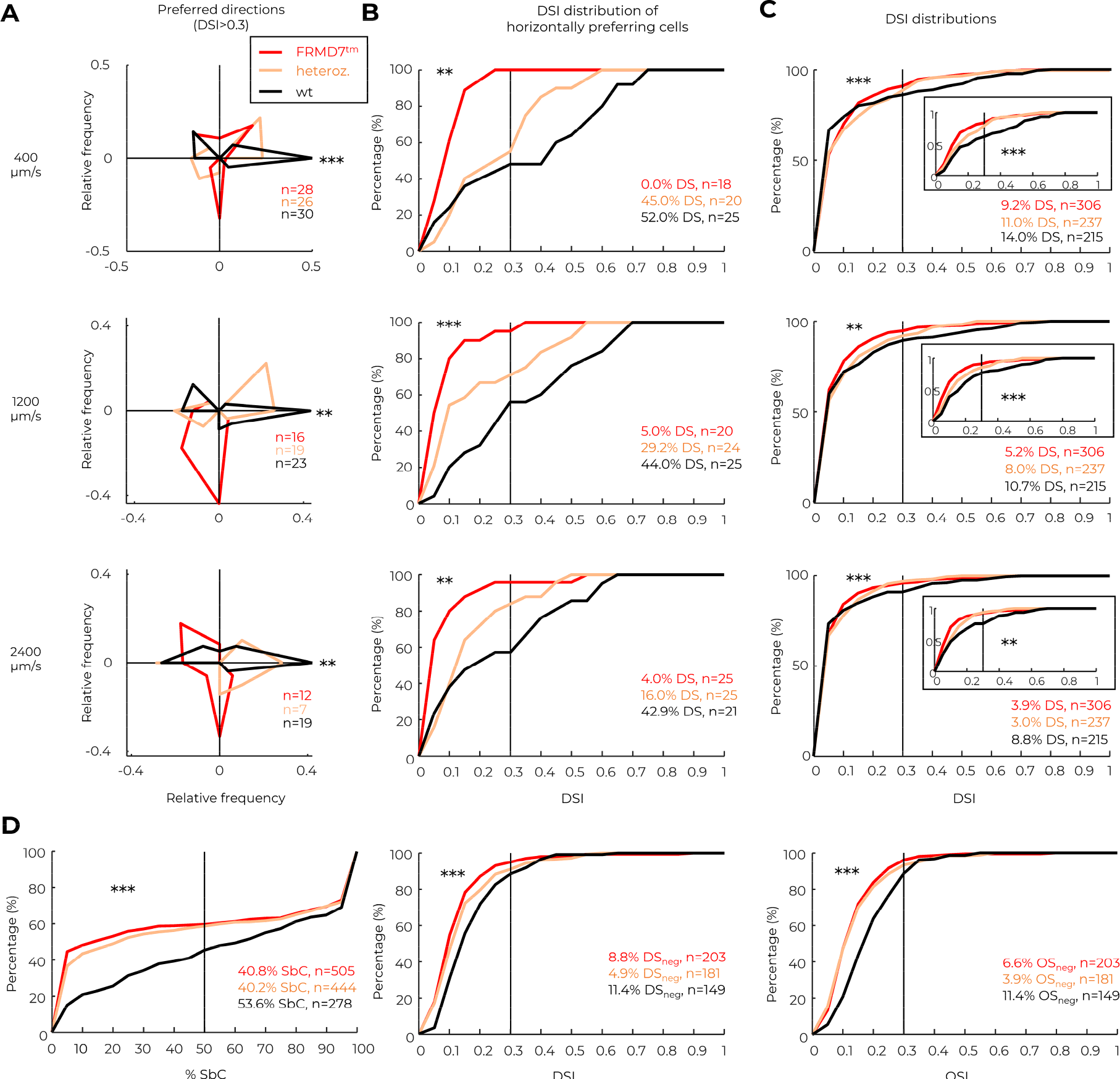
Horizontal direction selectivity is inherited from the retina. **A:** Polar plots of preferred directions of interneurons with DSI>0.3 in wild-type (black), hemizygous FRMD7^tm^ (red), and heterozygous FRMD7^tm^ (orange) mice. ***: p<0.001, **: p=0.007 (middle) and p=0.009 (bottom), Fisher’s exact test with Bonferroni-Holm correction. **B:** Cumulative histogram of DSI values across interneurons preferring horizontal motion (population vector direction 0 or 180° ± 15°) in wild-type (black), hemizygous FRMD7^tm^ (red), and heterozygous FRMD7^tm^ (orange) mice. ***: p<0.001, **: p=0.002 (upper panel), p=0.009 (lower panel), Kolmogorov-Smirnov test with Bonferroni-Holm correction. **C:** Cumulative histogram of DSI values across all interneurons in wild-type (black), hemizygous FRMD7^tm^ (red), and heterozygous FRMD7^tm^ (orange) mice. Inset: Distribution of DSI values across all interneurons for which the positive maximum response was larger than the negative maximum response. ***: p<0.001, **: p=0.005, Kolmogorov-Smirnov test with Bonferroni-Holm correction. **D**: Cumulative histogram of suppressed-by-contrast features in wild-type (black), hemizygous FRMD7^tm^ (red), and heterozygous FRMD7^tm^ (orange) mice. Left panel: Cumulative histogram of the percentage of responses suppressed by contrast. Middle/right panel: Cumulative histograms of the negative direction selectivity index (middle, DSI_neg_) and orientation selectivity index (right, OSI_neg_) for interneurons with >50% suppressed-by-contrast responses. ***: p<0.001, Kolmogorov-Smirnov test with Bonferroni-Holm correction. **A-D**: p-values refer to the comparison between hemizygous FRMD7^tm^ and wild-type mice. Interneurons recorded in 8 wild-type, 5 hemizygous FRMD7^tm^ und 3 heterozygous FRMD7^tm^ mice.

**Supplemental Figure 5:**
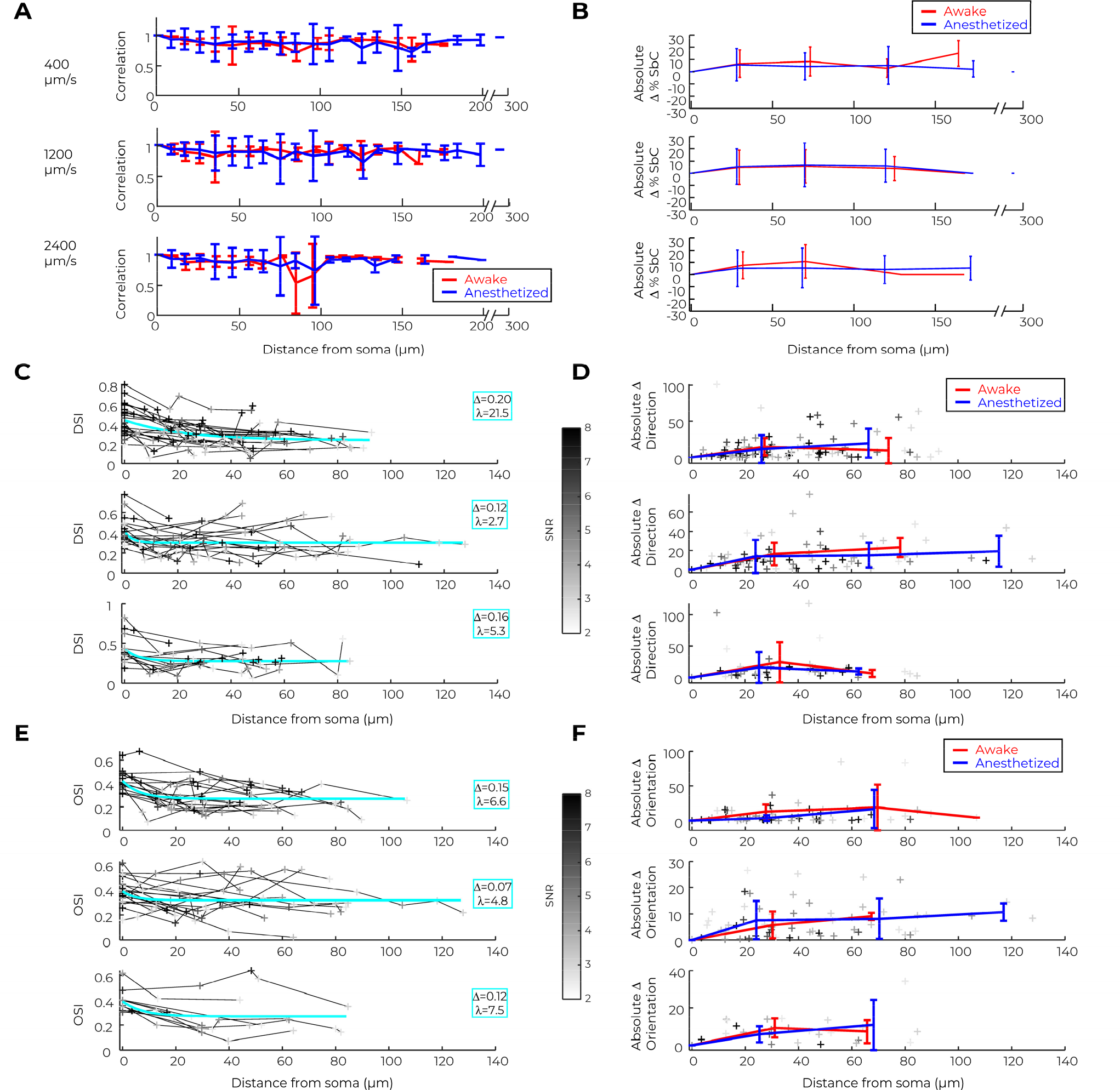
Somatic visual features extend into the dendrites. **A:** Correlation of the 16-dimensional response vectors (positive and negative response amplitudes to 8 directions) with their respective somatic response vector, plotted against Euclidean distance from the soma. Data averaged over 10µm bins. Blue: anesthetized, red: awake mice. Here and in all subsequent plots of **A**-**F**: Top: 400 µm/s; middle: 1200 µm/s; bottom: 2400 µm/s stimulus velocity. **B:** Absolute difference in the percentage of suppressed-by-contrast responses compared to the somatic responses, plotted against Euclidean distance from the soma. **C:** DSI values of individual compartments, plotted against Euclidean distance from the soma. **D:** Absolute difference in the preferred direction compared to the somatic response for data in **C**, plotted against Euclidean distance from the soma **E:** OSI values of individual compartments, plotted against Euclidean distance from the soma. **F:** Absolute difference in the preferred orientation compared to the somatic response for data in **E**, plotted against Euclidean distance from the soma **C, D**: Only cells with at least one compartment with DSI>0.3 included. **E, F**: Only cells with at least one compartment with OSI>0.3 included. **C, E**: Cyan: mono-exponential fit with length constant λ and amplitude Δ as indicated (y = Δ·exp(-x/λ) + y_0_). **B, D, F:** Red/blue: mean ± standard deviation of data recorded under awake (red) or anesthetized (blue) conditions in 50µm bins. Individual data points shaded in gray according to their signal-to-noise ratio (SNR). **A-F :** Interneurons recorded in 5 wild-type, 5 hemizygous FRMD7^tm^ und 3 heterozygous FRMD7^tm^ mice.

**Supplemental Figure 6:**
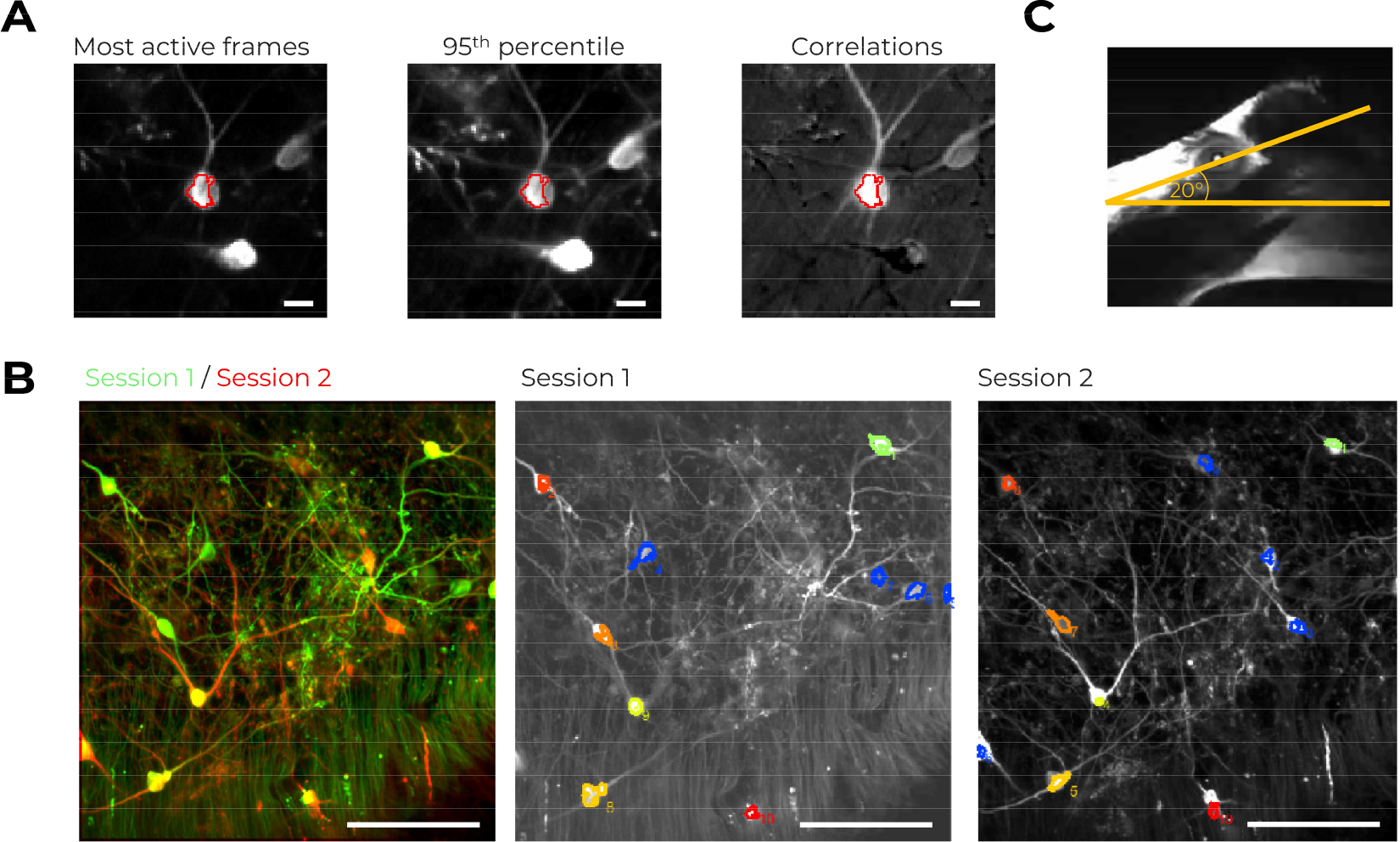
Method details of two-photon calcium imaging in vivo. **A: *Identification of interneuron somata*:** After selecting region of interest (ROI, red) with putative somata, the projection over the 500 most active frames (left panel), local 95% percentile projection (middle panel), and local correlation matrix (right panel) were inspected to confirm that the ROI corresponds to a soma. Scale bar: 10 µm. **B: *Avoiding duplication of cells*:** Recordings from the same or overlapping regions were aligned (left panel) and ROIs belonging to the same cells identified (right panels, ROIs belonging to the same cell plotted in the same color, non-matching ROIs in blue). Scale bar: 100 µm. **C: *Correction for head-mounting angle*:** The angle between the horizontal axis connecting the two eye corners and the table on which the stimulation screen was placed was determined. All stimulus directions were adjusted accordingly such that horizontal motion refers to motion along the horizontal axis of the eye.

## REFERENCES

1. Acuna-Goycolea, C., Brenowitz, S.D., Regehr, W.G., 2008. Active Dendritic Conductances Dynamically Regulate GABA Release from Thalamic Interneurons. Neuron 57, 420–431. https://doi.org/10.1016/j.neuron.2007.12.022

2. Baden, T., Berens, P., Bethge, M., Euler, T., 2013. Spikes in Mammalian Bipolar Cells Support Temporal Layering of the Inner Retina. Curr. Biol. 23, 48–52. https://doi.org/10.1016/j.cub.2012.11.006

3. Baden, T., Berens, P., Franke, K., Román Rosón, M., Bethge, M., Euler, T., 2016. The functional diversity of retinal ganglion cells in the mouse. Nature 529, 345–350. https://doi.org/10.1038/nature16468

4. Bae, J.A., Mu, S., Kim, J.S., Turner, N.L., Tartavull, I., Kemnitz, N., Jordan, C.S., Norton, A.D., Silversmith, W.M., Prentki, R., Sorek, M., David, C., Jones, D.L., Bland, D., Sterling, A.L.R., Park, J., Briggman, K.L., Seung, H.S., 2018. Digital Museum of Retinal Ganglion Cells with Dense Anatomy and Physiology. Cell 173, 1293–1306.e19. https://doi.org/10.1016/j.cell.2018.04.040

5. Bakken, T.E., van Velthoven, C.T., Menon, V., Hodge, R.D., Yao, Z., Nguyen, T.N., Graybuck, L.T., Horwitz, G.D., Bertagnolli, D., Goldy, J., Yanny, A.M., Garren, E., Parry, S., Casper, T., Shehata, S.I., Barkan, E.R., Szafer, A., Levi, B.P., Dee, N., Smith, K.A., Sunkin, S.M., Bernard, A., Phillips, J., Hawrylycz, M.J., Koch, C., Murphy, G.J., Lein, E., Zeng, H., Tasic, B., 2021. Single-cell and single-nucleus RNA-seq uncovers shared and distinct axes of variation in dorsal LGN neurons in mice, non-human primates, and humans. eLife 10, e64875. https://doi.org/10.7554/eLife.64875

6. Bauer, J., Weiler, S., Fernholz, M.H.P., Laubender, D., Scheuss, V., Hübener, M., Bonhoeffer, T., Rose, T., 2021. Limited functional convergence of eye-specific inputs in the retinogeniculate pathway of the mouse. Neuron 109, 2457–2468.e12. https://doi.org/10.1016/j.neuron.2021.05.036

7. Bickford, M.E., Zhou, N., Krahe, T.E., Govindaiah, G., Guido, W., 2015. Retinal and Tectal “Driver-Like” Inputs Converge in the Shell of the Mouse Dorsal Lateral Geniculate Nucleus. J. Neurosci. 35, 10523–10534. https://doi.org/10.1523/JNEUROSCI.3375-14.2015

8. Blitz, D.M., Regehr, W.G., 2005. Timing and Specificity of Feed-Forward Inhibition within the LGN. Neuron 45, 917–928. https://doi.org/10.1016/j.neuron.2005.01.033

9. Bloomfield, S.A., Hamos, J.E., Sherman, S.M., 1987. Passive cable properties and morphological correlates of neurones in the lateral geniculate nucleus of the cat. J. Physiol. 383, 653–692. https://doi.org/10.1113/jphysiol.1987.sp016435

10. Bloomfield, S.A., Sherman, S.M., 1989. Dendritic current flow in relay cells and interneurons of the cat’s lateral geniculate nucleus. Proc. Natl. Acad. Sci. 86, 3911–3914. https://doi.org/10.1073/pnas.86.10.3911

11. Casale, A.E., McCormick, D.A., 2011. Active Action Potential Propagation But Not Initiation in Thalamic Interneuron Dendrites. J. Neurosci. 31, 18289–18302. https://doi.org/10.1523/JNEUROSCI.4417-11.2011

12. Cleland, B.G., Dubin, M.W., Levick, W.R., 1971. Simultaneous Recording of Input and Output of Lateral Geniculate Neurones. Nature. New Biol. 231, 191–192. https://doi.org/10.1038/newbio231191a0

13. Colonnier, M., Guillery, R.W., 1964. Synaptic organization in the lateral geniculate nucleus of the monkey. Z. Für Zellforsch. Mikrosk. Anat. 62, 333–355. https://doi.org/10.1007/BF00339284

14. Cox, C.L., Sherman, S.M., 2000. Control of Dendritic Outputs of Inhibitory Interneurons in the Lateral Geniculate Nucleus. Neuron 27, 597–610. https://doi.org/10.1016/S0896-6273(00)00069-6

15. Cox, C.L., Zhou, Q., Sherman, S.M., 1998. Glutamate locally activates dendritic outputs of thalamic interneurons. Nature 394, 478–482. https://doi.org/10.1038/28855

16. Crandall, S.R., Cox, C.L., 2013. Thalamic microcircuits: presynaptic dendrites form two feedforward inhibitory pathways in thalamus. J. Neurophysiol. 110, 470–480. https://doi.org/10.1152/jn.00559.2012

17. Cruz-Martín, A., El-Danaf, R.N., Osakada, F., Sriram, B., Dhande, O.S., Nguyen, P.L., Callaway, E.M., Ghosh, A., Huberman, A.D., 2014. A dedicated circuit links direction-selective retinal ganglion cells to the primary visual cortex. Nature 507, 358–361. https://doi.org/10.1038/nature12989

18. Dubin, M.W., Cleland, B.G., 1977. Organization of visual inputs to interneurons of lateral geniculate nucleus of the cat. J. Neurophysiol. 40, 410–427. https://doi.org/10.1152/jn.1977.40.2.410

19. Gabbott, P.L.A., Bacon, S.J., 1995. Co-localisation of NADPH diaphorase activity and GABA immunoreactivity in local circuit neurones in the medial prefrontal cortex (mPFC) of the rat. Brain Res. 699, 321–328. https://doi.org/10.1016/0006-8993(95)01084-9

20. Goetz, J., Jessen, Z.F., Jacobi, A., Mani, A., Cooler, S., Greer, D., Kadri, S., Segal, J., Shekhar, K., Sanes, J.R., Schwartz, G.W., 2022. Unified classification of mouse retinal ganglion cells using function, morphology, and gene expression. Cell Rep. 40, 111040. https://doi.org/10.1016/j.celrep.2022.111040

21. Govindaiah, G., Cox, C.L., 2006. Metabotropic Glutamate Receptors Differentially Regulate GABAergic Inhibition in Thalamus. J. Neurosci. 26, 13443–13453. https://doi.org/10.1523/JNEUROSCI.3578-06.2006

22. Grubb, M.S., Thompson, I.D., 2004. Biochemical and anatomical subdivision of the dorsal lateral geniculate nucleus in normal mice and in mice lacking the β2 subunit of the nicotinic acetylcholine receptor. Vision Res., The Mouse Visual System: From Photoreceptors to Cortex 44, 3365–3376. https://doi.org/10.1016/j.visres.2004.09.003

23. Guillery, R.W., 1969. The organization of synaptic interconnections in the laminae of the dorsal lateral geniculate nucleus of the cat. Z. Für Zellforsch. Mikrosk. Anat. 96, 1–38. https://doi.org/10.1007/BF00321474

24. Hammer, S., Monavarfeshani, A., Lemon, T., Su, J., Fox, M.A., 2015. Multiple Retinal Axons Converge onto Relay Cells in the Adult Mouse Thalamus. Cell Rep. 12, 1575–1583. https://doi.org/10.1016/j.celrep.2015.08.003

25. Hamos, J.E., Van Horn, S.C., Raczkowski, D., Uhlrich, D.J., Sherman, S.M., 1985. Synaptic connectivity of a local circuit neurone in lateral geniculate nucleus of the cat. Nature 317, 618– 621. https://doi.org/10.1038/317618a0

26. Hirsch, J.A., Wang, X., Sommer, F.T., Martinez, L.M., 2015. How Inhibitory Circuits in the Thalamus Serve Vision. Annu. Rev. Neurosci. 38, 309–329. https://doi.org/10.1146/annurev-neuro-071013-014229

27. Howarth, M., Walmsley, L., Brown, T.M., 2014. Binocular Integration in the Mouse Lateral Geniculate Nuclei. Curr. Biol. 24, 1241–1247. https://doi.org/10.1016/j.cub.2014.04.014

28. Jaepel, J., Hübener, M., Bonhoeffer, T., Rose, T., 2017. Lateral geniculate neurons projecting to primary visual cortex show ocular dominance plasticity in adult mice. Nat. Neurosci. 20, 1708– 1714. https://doi.org/10.1038/s41593-017-0021-0

29. Jiang, Q., Litvina, E.Y., Acarón Ledesma, H., Shu, G., Sonoda, T., Wei, W., Chen, C., 2022. Functional convergence of on-off direction-selective ganglion cells in the visual thalamus. Curr. Biol. 32, 3110–3120.e6. https://doi.org/10.1016/j.cub.2022.06.023

30. Kay, J.N., Huerta, I.D. la, Kim, I.-J., Zhang, Y., Yamagata, M., Chu, M.W., Meister, M., Sanes, J.R., 2011. Retinal Ganglion Cells with Distinct Directional Preferences Differ in Molecular Identity, Structure, and Central Projections. J. Neurosci. 31, 7753–7762. https://doi.org/10.1523/JNEUROSCI.0907-11.2011

31. Kim, E.J., Jacobs, M.W., Ito-Cole, T., Callaway, E.M., 2016. Improved Monosynaptic Neural Circuit Tracing Using Engineered Rabies Virus Glycoproteins. Cell Rep. 15, 692–699. https://doi.org/10.1016/j.celrep.2016.03.067

32. Kim, I.-J., Zhang, Y., Yamagata, M., Meister, M., Sanes, J.R., 2008. Molecular identification of a retinal cell type that responds to upward motion. Nature 452, 478–482. https://doi.org/10.1038/nature06739

33. Krahe, T.E., El-Danaf, R.N., Dilger, E.K., Henderson, S.C., Guido, W., 2011. Morphologically Distinct Classes of Relay Cells Exhibit Regional Preferences in the Dorsal Lateral Geniculate Nucleus of the Mouse. J. Neurosci. 31, 17437–17448. https://doi.org/10.1523/JNEUROSCI.4370-11.2011

34. Leist, M., Datunashvilli, M., Kanyshkova, T., Zobeiri, M., Aissaoui, A., Cerina, M., Romanelli, M.N., Pape, H.-C., Budde, T., 2016. Two types of interneurons in the mouse lateral geniculate nucleus are characterized by different h-current density. Sci. Rep. 6, 24904. https://doi.org/10.1038/srep24904

35. Liang, L., Fratzl, A., Goldey, G., Ramesh, R.N., Sugden, A.U., Morgan, J.L., Chen, C., Andermann, M.L., 2018. A Fine-Scale Functional Logic to Convergence from Retina to Thalamus. Cell 173, 1343–1355.e24. https://doi.org/10.1016/j.cell.2018.04.041

36. Ling, S., Pratte, M.S., Tong, F., 2015. Attention alters orientation processing in the human lateral geniculate nucleus. Nat. Neurosci. 18, 496–498. https://doi.org/10.1038/nn.3967

37. Litvina, E.Y., Chen, C., 2017. Functional Convergence at the Retinogeniculate Synapse. Neuron 96, 330–338.e5. https://doi.org/10.1016/j.neuron.2017.09.037

38. Madisen, L., Zwingman, T.A., Sunkin, S.M., Oh, S.W., Zariwala, H.A., Gu, H., Ng, L.L., Palmiter, R.D., Hawrylycz, M.J., Jones, A.R., Lein, E.S., Zeng, H., 2010. A robust and high-throughput Cre reporting and characterization system for the whole mouse brain. Nat. Neurosci. 13, 133–140. https://doi.org/10.1038/nn.2467

39. Marshel, J.H., Kaye, A.P., Nauhaus, I., Callaway, E.M., 2012. Anterior-Posterior Direction Opponency in the Superficial Mouse Lateral Geniculate Nucleus. Neuron 76, 713–720. https://doi.org/10.1016/j.neuron.2012.09.021

40. Martersteck, E.M., Hirokawa, K.E., Evarts, M., Bernard, A., Duan, X., Li, Y., Ng, L., Oh, S.W., Ouellette, B., Royall, J.J., Stoecklin, M., Wang, Q., Zeng, H., Sanes, J.R., Harris, J.A., 2017. Diverse Central Projection Patterns of Retinal Ganglion Cells. Cell Rep. 18, 2058–2072. https://doi.org/10.1016/j.celrep.2017.01.075

41. Masland, R.H., 2012. The Neuronal Organization of the Retina. Neuron 76, 266–280. https://doi.org/10.1016/j.neuron.2012.10.002

42. Mastronarde, D.N., 1992. Nonlagged relay cells and interneurons in the cat lateral geniculate nucleus: Receptive-field properties and retinal inputs. Vis. Neurosci. 8, 407–441. https://doi.org/10.1017/S0952523800004934

43. Miyamichi, K., Amat, F., Moussavi, F., Wang, C., Wickersham, I., Wall, N.R., Taniguchi, H., Tasic, B., Huang, Z.J., He, Z., Callaway, E.M., Horowitz, M.A., Luo, L., 2011. Cortical representations of olfactory input by trans-synaptic tracing. Nature 472, 191–196. https://doi.org/10.1038/nature09714

44. Montero, V.M., Zempel, J., 1985. Evidence for two types of GABA-containing interneurons in the A-laminae of the cat lateral geniculate nucleus: a double-label HRP and GABA- immunocytochemical study. Exp. Brain Res. 60, 603–609. https://doi.org/10.1007/BF00236949

45. Morgan, J.L., Berger, D.R., Wetzel, A.W., Lichtman, J.W., 2016. The Fuzzy Logic of Network Connectivity in Mouse Visual Thalamus. Cell 165, 192–206. https://doi.org/10.1016/j.cell.2016.02.033

46. Morgan, J.L., Lichtman, J.W., 2020. An Individual Interneuron Participates in Many Kinds of Inhibition and Innervates Much of the Mouse Visual Thalamus. Neuron 106, 468–481.e2. https://doi.org/10.1016/j.neuron.2020.02.001

47. Okigawa, S., Yamaguchi, M., Ito, K.N., Takeuchi, R.F., Morimoto, N., Osakada, F., 2021. Cell type- and layer-specific convergence in core and shell neurons of the dorsal lateral geniculate nucleus. J. Comp. Neurol. 529, 2099–2124. https://doi.org/10.1002/cne.25075

48. Pare, D., Dossi, R.C., Steriade, M., 1991. Three types of inhibitory postsynaptic potentials generated by interneurons in the anterior thalamic complex of cat. J. Neurophysiol. 66, 1190– 1204. https://doi.org/10.1152/jn.1991.66.4.1190

49. Peng, Y.-R., Tran, N.M., Krishnaswamy, A., Kostadinov, D., Martersteck, E.M., Sanes, J.R., 2017. Satb1 Regulates Contactin 5 to Pattern Dendrites of a Mammalian Retinal Ganglion Cell. Neuron 95, 869–883.e6. https://doi.org/10.1016/j.neuron.2017.07.019

50. Piscopo, D.M., El-Danaf, R.N., Huberman, A.D., Niell, C.M., 2013. Diverse visual features encoded in mouse lateral geniculate nucleus. J. Neurosci. Off. J. Soc. Neurosci. 33, 4642–4656. https://doi.org/10.1523/JNEUROSCI.5187-12.2013

51. Raczkowski, D., Hamos, J.E., Sherman, S.M., 1988. Synaptic circuitry of physiologically identified W-cells in the cat’s dorsal lateral geniculate nucleus. J. Neurosci. 8, 31–48. https://doi.org/10.1523/JNEUROSCI.08-01-00031.1988

52. Reese, B.E., 1988. ‘Hidden lamination’ in the dorsal lateral geniculate nucleus: the functional organization of this thalamic region in the rat. Brain Res. Rev. 13, 119–137. https://doi.org/10.1016/0165-0173(88)90017-3

53. Rivlin-Etzion, M., Zhou, K., Wei, W., Elstrott, J., Nguyen, P.L., Barres, B.A., Huberman, A.D., Feller, M.B., 2011. Transgenic Mice Reveal Unexpected Diversity of On-Off Direction-Selective Retinal Ganglion Cell Subtypes and Brain Structures Involved in Motion Processing. J. Neurosci. 31, 8760–8769. https://doi.org/10.1523/JNEUROSCI.0564-11.2011

54. Román Rosón, M., Bauer, Y., Kotkat, A.H., Berens, P., Euler, T., Busse, L., 2019. Mouse dLGN Receives Functional Input from a Diverse Population of Retinal Ganglion Cells with Limited Convergence. Neuron 102, 462–476.e8. https://doi.org/10.1016/j.neuron.2019.01.040

55. Rompani, S.B., Müllner, F.E., Wanner, A., Zhang, C., Roth, C.N., Yonehara, K., Roska, B., 2017. Different Modes of Visual Integration in the Lateral Geniculate Nucleus Revealed by Single-Cell-Initiated Transsynaptic Tracing. Neuron 93, 767–776.e6. https://doi.org/10.1016/j.neuron.2017.01.028

56. Ronneberger, O., Fischer, P., Brox, T., 2015. U-Net: Convolutional Networks for Biomedical Image Segmentation. https://doi.org/10.48550/arXiv.1505.04597

57. Roska, B., Werblin, F., 2001. Vertical interactions across ten parallel, stacked representations in the mammalian retina. Nature 410, 583–587. https://doi.org/10.1038/35069068

58. Sabbah, S., Gemmer, J.A., Bhatia-Lin, A., Manoff, G., Castro, G., Siegel, J.K., Jeffery, N., Berson, D.M., 2017. A retinal code for motion along the gravitational and body axes. Nature 546, 492–497. https://doi.org/10.1038/nature22818

59. Seabrook, T.A., Krahe, T.E., Govindaiah, G., Guido, W., 2013. Interneurons in the mouse visual thalamus maintain a high degree of retinal convergence throughout postnatal development. Neural Develop. 8, 24. https://doi.org/10.1186/1749-8104-8-24

60. Sherman, S.M., 2004. Interneurons and triadic circuitry of the thalamus. Trends Neurosci. 27, 670–675. https://doi.org/10.1016/j.tins.2004.08.003

61. Sherman, S.M., Friedlander, M.J., 1988. Identification of X versus Y properties for interneurons in the A-laminae of the cat’s lateral geniculate nucleus. Exp. Brain Res. 73, 384–392. https://doi.org/10.1007/BF00248231

62. Sherman, S.M., Guillery, R.W., 2009. Exploring the Thalamus and Its Role in Cortical Function. https://doi.org/10.7551/mitpress/2940.001.0001

63. Siegert, S., Scherf, B.G., Del Punta, K., Didkovsky, N., Heintz, N., Roska, B., 2009. Genetic address book for retinal cell types. Nat. Neurosci. 12, 1197–1204. https://doi.org/10.1038/nn.2370

64. Sommeijer, J.-P., Ahmadlou, M., Saiepour, M.H., Seignette, K., Min, R., Heimel, J.A., Levelt, C.N., 2017. Thalamic inhibition regulates critical-period plasticity in visual cortex and thalamus. Nat. Neurosci. 20, 1715–1721. https://doi.org/10.1038/s41593-017-0002-3

65. Szentágothai, J., Hámori, J., Tömböl, T., 1966. Degeneration and electron microscope analysis of the synaptic glomeruli in the lateral geniculate body. Exp. Brain Res. 2, 283–301. https://doi.org/10.1007/BF00234775

66. Tamamaki, N., Yanagawa, Y., Tomioka, R., Miyazaki, J.-I., Obata, K., Kaneko, T., 2003. Green fluorescent protein expression and colocalization with calretinin, parvalbumin, and somatostatin in the GAD67-GFP knock-in mouse. J. Comp. Neurol. 467, 60–79. https://doi.org/10.1002/cne.10905

67. Taniguchi, H., He, M., Wu, P., Kim, S., Paik, R., Sugino, K., Kvitsani, D., Fu, Y., Lu, J., Lin, Y., Miyoshi, G., Shima, Y., Fishell, G., Nelson, S.B., Huang, Z.J., 2011. A Resource of Cre Driver Lines for Genetic Targeting of GABAergic Neurons in Cerebral Cortex. Neuron 71, 995–1013. https://doi.org/10.1016/j.neuron.2011.07.026

68. Tran, N.M., Shekhar, K., Whitney, I.E., Jacobi, A., Benhar, I., Hong, G., Yan, W., Adiconis, X., Arnold, M.E., Lee, J.M., Levin, J.Z., Lin, D., Wang, C., Lieber, C.M., Regev, A., He, Z., Sanes, J.R., 2019. Single-Cell Profiles of Retinal Ganglion Cells Differing in Resilience to Injury Reveal Neuroprotective Genes. Neuron 104, 1039–1055.e12. https://doi.org/10.1016/j.neuron.2019.11.006

69. Usrey, W.M., Reppas, J.B., Reid, R.C., 1999. Specificity and Strength of Retinogeniculate Connections. J. Neurophysiol. 82, 3527–3540. https://doi.org/10.1152/jn.1999.82.6.3527

70. Vigeland, L.E., Contreras, D., Palmer, L.A., 2013. Synaptic Mechanisms of Temporal Diversity in the Lateral Geniculate Nucleus of the Thalamus. J. Neurosci. 33, 1887–1896. https://doi.org/10.1523/JNEUROSCI.4046-12.2013

71. Wang, X., Sommer, F.T., Hirsch, J.A., 2011. Inhibitory circuits for visual processing in thalamus. Curr. Opin. Neurobiol., Networks, circuits and computation 21, 726–733. https://doi.org/10.1016/j.conb.2011.06.004

72. Wässle, H., 2004. Parallel processing in the mammalian retina. Nat. Rev. Neurosci. 5, 747–757. https://doi.org/10.1038/nrn1497

73. Wertz, A., Trenholm, S., Yonehara, K., Hillier, D., Raics, Z., Leinweber, M., Szalay, G., Ghanem, A., Keller, G., Rózsa, B., Conzelmann, K.-K., Roska, B., 2015. PRESYNAPTIC NETWORKS. Single-cell-initiated monosynaptic tracing reveals layer-specific cortical network modules. Science 349, 70–74. https://doi.org/10.1126/science.aab1687

74. Wickersham, I.R., Lyon, D.C., Barnard, R.J.O., Mori, T., Finke, S., Conzelmann, K.-K., Young, J.A.T., Callaway, E.M., 2007. Monosynaptic Restriction of Transsynaptic Tracing from Single, Genetically Targeted Neurons. Neuron 53, 639–647. https://doi.org/10.1016/j.neuron.2007.01.033

75. Wienbar, S., Schwartz, G.W., 2022. Differences in spike generation instead of synaptic inputs determine the feature selectivity of two retinal cell types. Neuron 110, 2110–2123.e4. https://doi.org/10.1016/j.neuron.2022.04.012

76. Wilson, J.R., Friedlander, M.J., Sherman, S.M., 1984. Fine structural morphology of identified X- and Y-cells in the cat’s lateral geniculate nucleus. Proc. R. Soc. Lond. B Biol. Sci. 221, 411–436. https://doi.org/10.1098/rspb.1984.0042

77. Yonehara, K., Fiscella, M., Drinnenberg, A., Esposti, F., Trenholm, S., Krol, J., Franke, F., Scherf, B.G., Kusnyerik, A., Müller, J., Szabo, A., Jüttner, J., Cordoba, F., Reddy, A.P., Németh, J., Nagy, Z.Z., Munier, F., Hierlemann, A., Roska, B., 2016. Congenital Nystagmus Gene FRMD7 Is Necessary for Establishing a Neuronal Circuit Asymmetry for Direction Selectivity. Neuron 89, 177–193. https://doi.org/10.1016/j.neuron.2015.11.032

78. Zeater, N., Cheong, S.K., Solomon, S.G., Dreher, B., Martin, P.R., 2015. Binocular Visual Responses in the Primate Lateral Geniculate Nucleus. Curr. Biol. 25, 3190–3195. https://doi.org/10.1016/j.cub.2015.10.033

79. Zhu, J.J., Uhlrich, D.J., Lytton, W.W., 1999. Properties of a hyperpolarization-activated cation current in interneurons in the rat lateral geniculate nucleus. Neuroscience 92, 445–457. https://doi.org/10.1016/S0306-4522(98)00759-3

